# Decomposing response inhibition: a POMDP model

**DOI:** 10.64898/2026.02.26.708416

**Authors:** Wenting Wang, Tobias Kaufmann, Peter Dayan

## Abstract

Inhibitory control is a core cognitive function whose competence varies across the population, with impairments often observed in psychiatric conditions such as attention deficit hyperactivity disorder (ADHD). The Stop Signal Task (SST) is a widely used paradigm for assessing this ability. However, conventional formalizations of SST performance, such as the independent race model, rely on assumptions that are frequently violated in modern experimental designs. Furthermore, they typically fit only mean reaction times, overlooking crucial trial-by-trial dynamics. To address these limitations, we formalize the SST as a partially observable Markov decision process (POMDP). This framework characterizes inhibitory control with two components: noisy perceptual inference regarding stimuli and optimal control balanced against potential costs. To fit this model to the Adolescent Brain Cognitive Development (ABCD) study baseline cohort (N = 3,567), we introduce Transformer-encoded Simulation-Based Inference (TeSBI). This end-to-end architecture learns compact, sequence-aware embeddings from raw behavioral data. It enables efficient, amortized inference of individual-level posteriors. Extensive validation confirms it extracts reliable and identifiable parameters. We identify distinct latent computational attributes associated with scores on ADHD questionnaires. Controlling for sex, IQ, and medication status, children with higher ADHD scores exhibit subtle but robust shifts in computational attributes. They show a reduction in go cue directional precision, alongside a blunted sensitivity to stop error, go error and time costs. The learned embedding space reveals a continuous manifold in which children with higher ADHD scores are heterogeneously distributed, rather than forming distinct disorder clusters. This indicates that similar clinical characteristics can emerge from diverse combinations of computational mechanisms, supporting a dimensional perspective on neurodiversity. Our end-to-end framework can be extended to a broader range of cognitive tasks. It offers a scalable, theory-driven solution for analyzing large-scale behavioral data.

**Author summary:** Inhibitory control is essential for adjusting thoughts and behavior and is often impaired in conditions like ADHD. Traditional assessments often oversimplify the underlying decision-making. We address this using a biologically grounded mathematical framework that separates how we perceive signals from how we execute control. This framework remains valid in diverse experimental designs where traditional models fail. To apply this complex model at scale, we develop a specialized machine learning approach (TeSBI) to reverse-engineer individual cognitive profiles. Analyzing data from over 3,000 children, we linked higher ADHD scores to subtle, consistent computational shifts: a reduction in go cue directional precision, a blunted sensitivity to the costs of making mistakes or taking too much time. These alterations remain robust regardless of sex, IQ, or medication status. Rather than forming a single disorder cluster, these children displayed diverse cognitive profiles, supporting a dimensional view of neurodiversity. By combining theory-driven cognitive modeling with scalable data-driven inference, our framework effectively captures complex inhibitory mechanisms and enables the precise analysis of large-scale behavioral datasets. This approach paves the way for more personalized strategies in computational psychiatry by recognizing the inherent heterogeneity within clinical characteristics.

## Introduction

Inhibitory control is a fundamental cognitive function that regulates thoughts, actions, and emotional responses. It is essential for flexible and goal-directed behavior in dynamic environments. This capacity is often impaired in psychiatric conditions such as attention deficit hyperactivity disorder (ADHD), autism spectrum disorder, and obsessive-compulsive disorder [1–3].

One popular method for examining individual differences in inhibitory control is the Stop Signal Task (SST) [4, 5]. In this task, participants must make a fast *go* response to a cue unless a subsequent stop signal countermands the action. The interval between the go cue and the stop signal is the Stop Signal Delay (SSD). A shorter SSD makes inhibition easier. The SSD is typically adjusted on a trial-by-trial basis using a tracking algorithm (a *staircase*) to maintain a fixed proportion of inhibition failures. Standard SSTs are commonly modeled using evidence accumulation frameworks, such as the race model [4–6] or the drift diffusion model [7, 8]. The parameters derived from these models provide significant insights into various group differences [9–12].

However, these standard models typically rely on subject-level aggregate metrics (e.g., mean reaction time, accuracy), which fail to capture the full complexity of behavioral dynamics. Furthermore, they assume that go and stop signals are processed independently. Neural evidence contradicts this assumption even in the standard SST [13, 14], and some modern experimental designs directly violate it. A prominent example is the SST in the Adolescent Brain Cognitive Development (ABCD) study [15], where the stop signal visually masks the preceding go cue. In such contexts, race models relying on an assumption that go and stop signals are independent lead to biased estimates of the Stop Signal Reaction Time (SSRT) [16, 17].

To address these limitations, we propose a general computational framework grounded in the Partially Observable Markov Decision Process (POMDP) formalism [18–23]. This framework naturally accommodates interactive go and stop processes while unifying perceptual inference and optimal control. By explicitly parameterizing key mechanisms, such as sensory precision, intrinsic costs, and probabilistic action selection, the POMDP model captures a wide range of behavioral variability across a heterogeneous population.

Fitting this model to the ABCD dataset presents computational challenges due to intractable likelihood functions, the adaptive SSD staircase, and sequential effects across trials. To overcome these obstacles, we introduce Transformer-encoded Simulation-Based Inference (TeSBI) [24–26]. Unlike traditional methods that rely on handcrafted summary statistics, the Transformer encoder learns compact, sequence-aware representations of behavioral data, preserving temporal dependencies and contextual information. A normalizing flow [27] enables an efficient and reliable estimation of individual-level parameter posteriors. Together, this end-to-end framework facilitates scalable model fitting for large-scale datasets.

In this study, we apply the POMDP-TeSBI framework to the ABCD SST dataset.

Following rigorous model selection and comprehensive validation, we characterize individual perceptual capabilities and intrinsic valuation. We report the resulting subject-level parameter distributions and their small, but significant, associations with scores on ADHD questionnaires. Finally, we visualize distinct cognitive profiles using the embedding space learned by the Transformer encoder.

## Materials and methods

### Ethics statement

Study protocols have been approved by either local institutional review boards (IRB) or by reliance agreements with the central IRB at the University of California [28].

Written informed consent was obtained from the parents or legal guardians of all child participants, and written assent was obtained from the children themselves, prior to participation [29].

### Experimental task and mental health measures

#### Stop signal task in the ABCD study

To examine inhibitory control, we analyze data from the Stop Signal Task (SST) deployed in the Adolescent Brain Cognitive Development (ABCD) Study, Annual Release 5.1 [15].

SST experimental designs are generally categorized as independent or dependent. In an independent design, the go cue remains visible when the stop signal appears. Classic formalizations, such as the independent race model [5], duly assume that go and stop signals are processed independently (although some neural evidence weighs against this assumption [13, 14]). In contrast, the ABCD study utilizes a dependent design (Fig 1). In this version, the stop signal visually replaces, or masks, the preceding go cue. This replacement would disrupt and slow the processing of the go signal specifically on stop trials. Consequently, the core race model assumption that the go process remains identical across trial types is violated. This structural dependence confounds standard SSRT calculations [17].

**Fig 1.**
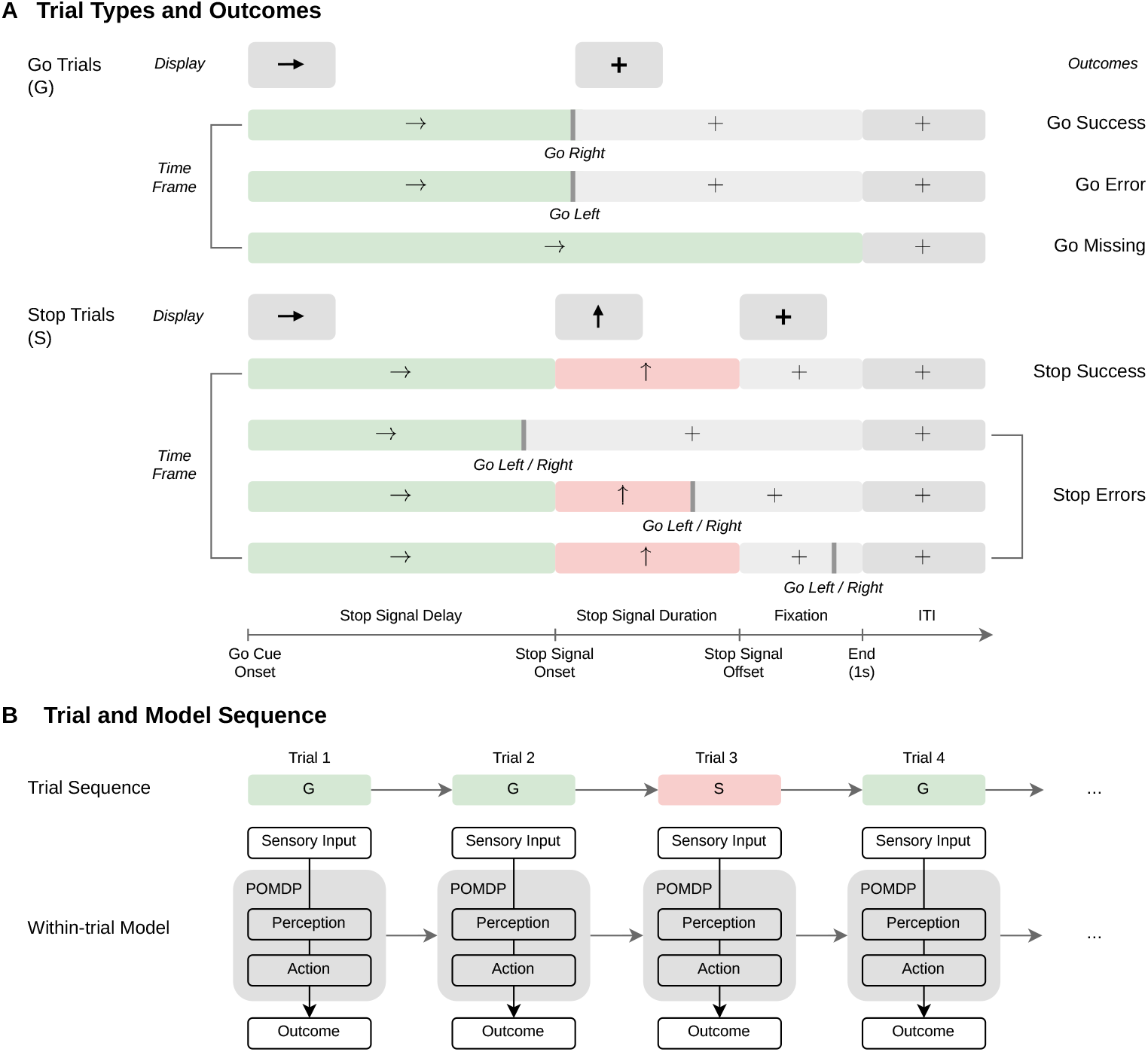
ABCD stop signal task and POMDP framework. (A) The task consists of 360 trials (300 go trials, 60 stop trials). Each trial has a fixed duration of 1000 ms. Black stimuli appear on a mid-gray background (top rectangles). Colored bars below illustrate the trial timing. When a response is registered, a fixation cross replaces the active stimulus for the remainder of the trial. On stop trials, a response registered at any point is categorized as a stop error. This includes responses during the initial Stop Signal Delay (SSD), the stop signal duration, or the subsequent fixation phase. The SSD is adjusted on a trial-by-trial basis to maintain a stop success rate of approximately 50%. (B) The POMDP framework models the within-trial dynamics of perception and choice. It is fit to the full behavioral dynamics, including the SSD staircase. The trial sequence shown is illustrative. Actual ABCD sessions follow predefined pseudo-randomized orders.

Participants were instructed to press a button corresponding to the direction of a go cue (a left or right arrow presented with equal probability) as quickly and accurately as possible. They were instructed to withhold this response if a stop signal (an upward-pointing arrow) subsequently appeared. The session consisted of 360 trials divided into two blocks of 180 trials each. One-sixth of these (60 trials) were stop trials.

All trials had a fixed duration of 1000 ms. They were separated by an inter-trial interval (ITI) jittered between 700 and 2000 ms. On go trials, the go cue remained on the screen until a response occurred or until a 1000-ms timeout was reached. Following a response, a fixation cross appeared for the remainder of the trial. On stop trials, the go cue appeared for a variable Stop Signal Delay (SSD). The stop signal then replaced the go cue for 300 ms, or until the 1000 ms trial limit if the SSD was *≥* 700 ms. If the participant responded despite the stop signal (i.e., failed inhibition), the stimulus was immediately replaced by a fixation cross.

The SSD was adjusted on a trial-by-trial basis using a tracking algorithm. This algorithm was designed to maintain a stop success rate of approximately 50%. The initial SSD was set to 50 ms. It increased by 50 ms following a successful inhibition and decreased by 50 ms following a failed inhibition. The SSD was intended to be constrained within a range of 50 to 850 ms.

We focused on baseline data from the ABCD study sample. We applied rigorous quality control criteria that evaluated both the technical experimental setup and participant task compliance. These inclusion criteria required: (1) at least three events for all trial types and outcomes; (2) no box switching due to reversed hardware mapping; (3) no task coding errors disrupting the SSD tracking algorithm; (4) fewer than four stop trials with a 0 ms SSD (these were typically caused by technical difficulties); (5) a mean reaction time (RT) on stop error trials not exceeding the mean RT on correct go trials; (6) a complete session of 360 trials, including exactly 60 stop trials; (7) fewer than four extremely fast trials (RT ≤200 ms); (8) a go missing rate ≤ 10%; (9) a go error rate ≤ 10%; and (10) no missing or incomplete demographic data.

After applying these exclusions, the final sample consisted of 3,567 participants. Further details regarding data quality control and model-agnostic statistical analyses are provided in Section F of S1 Appendix.

#### Mental health measures

At the baseline year (children aged 9-10 years), ADHD symptomatology was assessed using the parent-reported Child Behavior Checklist (CBCL). We utilized the DSM-5 ADHD raw scores [30]. These scores range from 0 to 14. Higher values indicate greater symptom severity. A continuous measure of ADHD symptomatology provides a more nuanced characterization of individual differences than binary diagnostic categories. This dimensional approach aligns with the National Institute of Mental Health’s Research Domain Criteria (RDoC) framework [31]. Further details regarding CBCL administration are available in the ABCD Study protocols.

### Model framework: POMDP

Consistent with established frameworks for perceptual decision-making [18, 21, 22, 32], we formulate the SST as a Partially Observable Markov Decision Process (POMDP). We model it using two coupled observational streams: one for the directional go cue and one for the inhibitory stop signal. These streams are integrated via Bayesian inference to generate evolving belief states regarding the trial direction (Left versus Right) and trial type (Go versus Stop).

At each time step, based on these beliefs, the agent selects an action (Go-Left, o-Right, or Wait) derived from a policy that optimizes total return. By balancing the costs of temporal delay, directional errors, missed responses, and failed inhibition, the model unifies Bayesian inference with optimal control. All model parameters are defined in Table 1 and detailed below.

**Table 1.** POMDP model parameters and descriptions Parameter Name Description.

| Parameter | Name | Description |
| --- | --- | --- |
| *Sensory Parameters* |  |  |
| $\chi'$ | Go null | Probability of receiving uninformative noise regarding the go cue (sensory lapse/ambiguity). |
| $\chi$ | Go precision | Probability of successfully resolving the go cue direction (sensory fidelity). |
| $\delta'$ | Stop null | Probability of receiving uninformative noise regarding the stop signal (sensory lapse/ambiguity). |
| $\delta$ | Stop precision | Probability of successfully resolving the presence or absence of the stop signal (signal fidelity). |
| *Cost Parameters* |  |  |
| $c_{ge}$ | Go error cost | Subjective penalty for committing a directional error on go trials. |
| $c_{gm}$ | Go missing cost | Subjective penalty for failing to respond (missing) on go trials. |
| $c_{se}$ | Stop error cost | Subjective penalty for failing to inhibit the response on stop trials. |
| $c_t$ | Time cost | Subjective penalty for each time step spent waiting. |
| *Control and Dynamic Parameters* |  |  |
| $\varphi$ | Inverse temperature | Degree of determinism in softmax policy. |
| $\lambda$ | Hazard rate | Temporal prior of stop signal onset. |
| $\tau$ | Non-decision time | Sensory transmission and motor execution delays. |
| $\tilde{r}^\nu$ | Stop prior | Predicted prior probability that the trial $\nu$ will be a stop trial. |

Our POMDP framework accommodates different SST experimental designs. The main text focuses on a *dependent* design such as the ABCD task. As noted, in this design, the stop signal visually masks the go cue, causing their sensory observation processes to interact. The model also generalizes to *independent* designs. In these designs, the go and stop stimuli remain simultaneously visible without perceptual interference. Specifications for modeling independent SSTs are detailed in Section A.2 of S1 Appendix.

### Perceptual inference

We follow standard Bayesian treatments of perceptual processing [22, 32]. On each trial *ν*, a generative process specifies how latent states produce observable sensory inputs at each time step *t*. The inference process then uses these observations to update beliefs about the latent states (Fig 2). We discretize the 1-second trial duration into *T* = 40 time steps of 25 ms each. For brevity, we omit the trial superscript *ν* where contextually clear.

**Fig 2.**
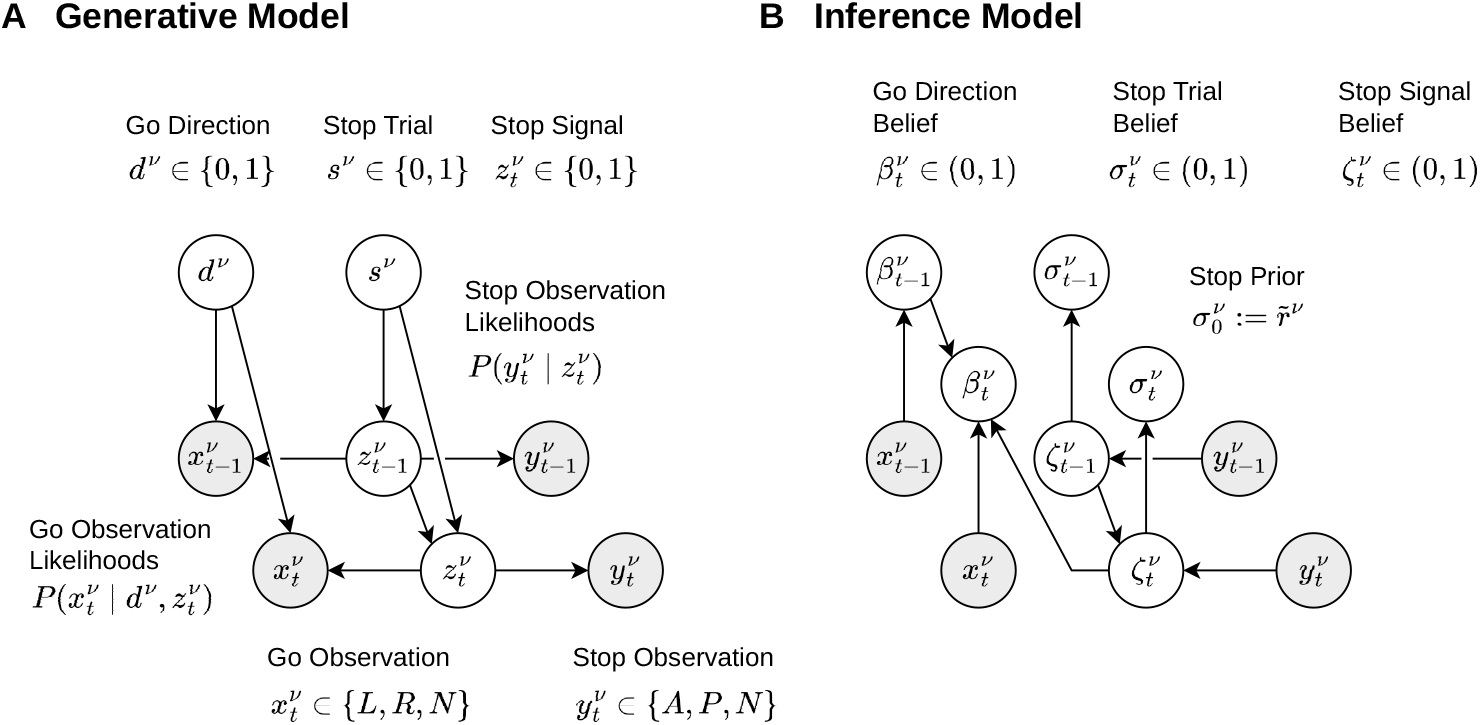
Generative and inference processes in the POMDP model. (A) Generative model. Each trial *ν* has a true go direction *d*^*ν*^ (Left or Right) and a true trial type *s*^*ν*^ (Go or Stop). At each time step *t, s*^*ν*^ and the Stop Signal Delay (SSD) determine the true stop signal state 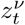 (Absence or Presence). The latent states *d*^*ν*^, *s*^*ν*^, and 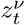 are hidden from the participant. Observable go cue inputs 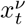 depend on both *d*^*ν*^ and 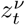. Observable stop signal inputs 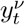 depend only on 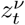. (B) Inference model. At each time step *t*, the participant uses Bayes rule to calculate belief states (probabilities of the latent states). The stop signal state 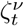 is inferred from 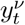 and a hazard function. The trial type belief 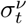 is derived directly from 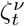.The go direction belief 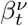 is inferred from 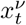 and 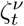.

#### The stop process: generative and inference models

In the generative model, the latent state *s*^*ν*^ ∈ {0, 1} indicates the true trial type (0 for Go trial, 1 for Stop trial). On stop trials, the stop signal appears at a specific stop signal delay (SSD, *θ*^*ν*^) and lasts for *µ* + 1 = 12 time steps (300 ms). The true stop signal state at time *t* is denoted as 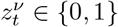 (0 for Absence, 1 for Presence). Participants receive a sensory observation 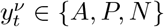 (Absence, Presence, or Null), generated from a multinomial distribution conditioned on 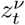. As detailed in Table 2, these observation likelihoods shift across the pre-signal, signal-on, and post-signal phases.

**Table 2.**
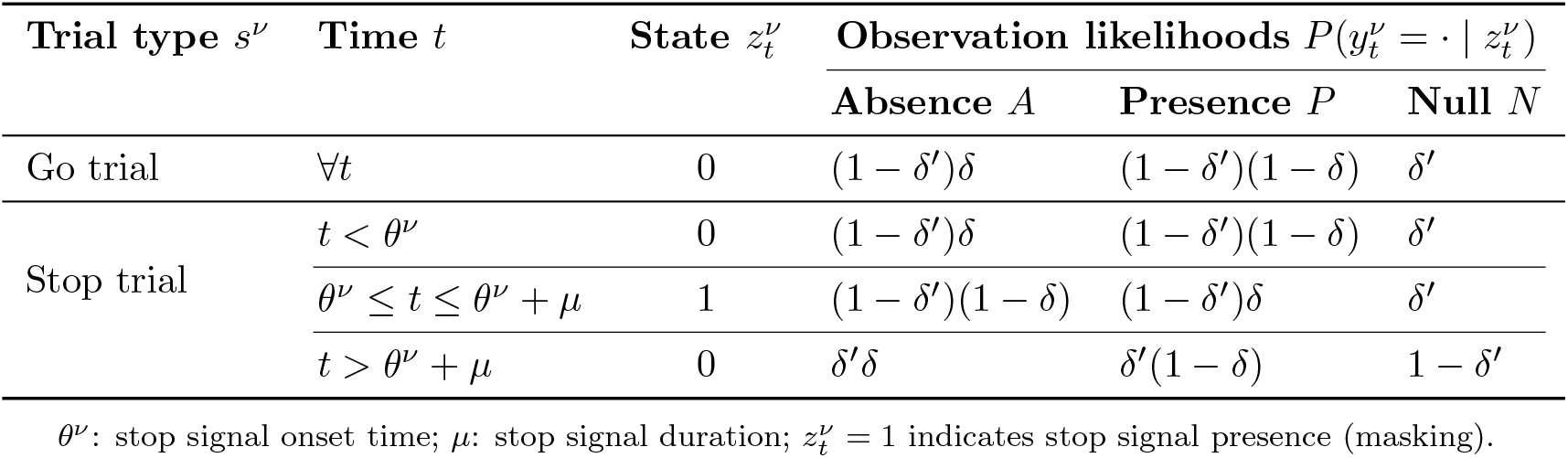
Observation likelihoods for the stop process.

The inference model estimates the posterior probability that the current trial is a stop trial, 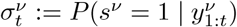. To compute this efficiently, the agent tracks an intermediate belief 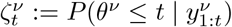 representing the cumulative probability that the stop signal has *already* occurred.

To align with participants’ behavioral constraints, we introduce three simplifying assumptions: (1) the stop belief is updated solely based on stop observations 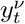 (2) participants expect the stop signal onset based on a constant hazard rate *λ*; and (3) participants model the transient stop signal as persisting until the trial ends (although the input statistics change according to the bottom row of table 2). This avoids complex inference about signal offset. It is justified empirically: only 10.03% of stop errors occurred during the post-signal fixation phase.

Based on these assumptions, the belief 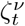 is updated recursively via Bayes rule. First, the predicted belief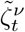 incorporates the hazard function for the occurrence of the stop signal, 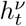, which combines the temporal prior *λ* and the stop prior 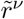:

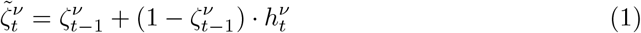

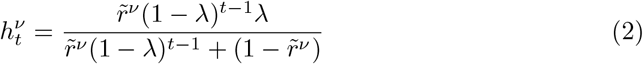

Next, the posterior 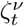 is corrected by the current sensory observation 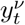:

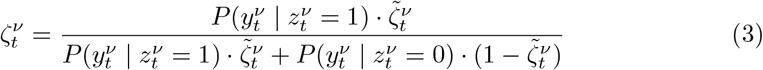

Finally, the total belief that the trial is a stop trial, 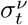, integrates 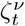 with the residual probability of a future onset:

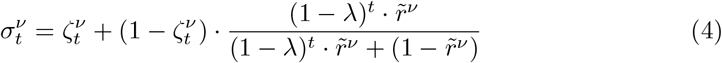

Detailed derivation of stop process inference is in Section A.1 of the S1 Appendix.

#### The go process: generative and inference models

In the generative model, the latent state *d*^*ν*^ ***∈*** {0, 1} denotes the true direction of the go cue (0 for Left, 1 for Right). Participants receive a noisy sensory input 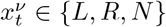 (Left, Right, or Null) at each time step. This input is governed by the go precision (*χ*) and go null (*χ*^*′*^) parameters (Table 3). On go trials, the cue remains visible throughout. On stop trials, the appearance of the stop signal at time *θ*^*ν*^ visually masks the go cue. This masking shifts the observation 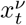 predominantly toward the Null status for the remainder of the trial (albeit allowing for a small residual signal associated with the go direction).

**Table 3.** Observation likelihoods for the go process.

| Trial type $s^\nu$ | Time $t$ | State $z_t^\nu$ | State $d^\nu$ | Observation likelihoods $P(x_t^\nu = \cdot \vert d^\nu, z_t^\nu)$ | | |
| --- | --- | --- | --- | --- | --- | --- |
| | | | | Left $L$ | Right $R$ | Null $N$ |
| Go trial | $\forall t$ | 0 | 0 | $(1 - \chi')\chi$ | $(1 - \chi')(1 - \chi)$ | $\chi'$ |
| | | | 1 | $(1 - \chi')(1 - \chi)$ | $(1 - \chi')\chi$ | $\chi'$ |
| Stop trial | $t < \theta^\nu$ | 0 | 0 | $(1 - \chi')\chi$ | $(1 - \chi')(1 - \chi)$ | $\chi'$ |
| | | | 1 | $(1 - \chi')(1 - \chi)$ | $(1 - \chi')\chi$ | $\chi'$ |
| | $t \geq \theta^\nu$ | 1 | 0 | $\chi'\chi$ | $\chi'(1 - \chi)$ | $1 - \chi'$ |
| | | | 1 | $\chi'(1 - \chi)$ | $\chi'\chi$ | $1 - \chi'$ |

$\theta^\nu$ : stop signal onset time; $\mu$ : stop signal duration; $z_t^\nu = 1$ indicates stop signal presence (masking).

The inference model estimates the directional belief state 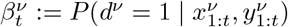 It is initialized uniformly at 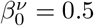 (the probability of a left cue is 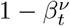). The presence of the stop signal 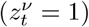 masks the go cue and alters its observation likelihoods. Therefore, the inference for *d*^*ν*^ depends partially on the current stop signal belief 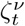.

The belief 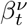 is updated recursively via Bayes rule by marginalizing over the stop signal state *z*:

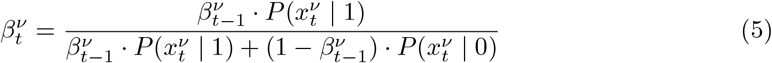

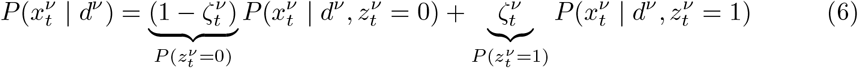

where 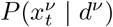 is the marginal likelihood of observing 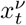 given direction *d*^*ν*^. An ambiguous sensory input 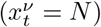 provides no directional evidence, resulting in no belief update 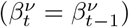.

The complete belief state at time *t* of trial *ν* is formalized as 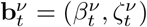.This joint state encodes the participant’s inferred probabilities regarding both the go direction and the stop signal state. The trial type probability 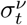 is deterministically derived from 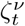 (Eq 4). Therefore, maintaining 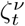 acts as a sufficient statistic for the stop process within the state representation.

### Optimal control

On each time step *t*, the participant chooses an action *a*_*t*_ **∈** {*L, R, W*} (Go-Left, Go-Right or Wait). We formulate the action-value *Q*_*t*_(**b**_*t*_, *a*_*t*_) in terms of expected *cost*, which is determined by the current belief state **b**_*t*_ = (*β*_*t*_, *ζ*_*t*_) and intrinsic cost parameters. The objective is to minimize the total cost (Fig 3).

**Fig 3.**
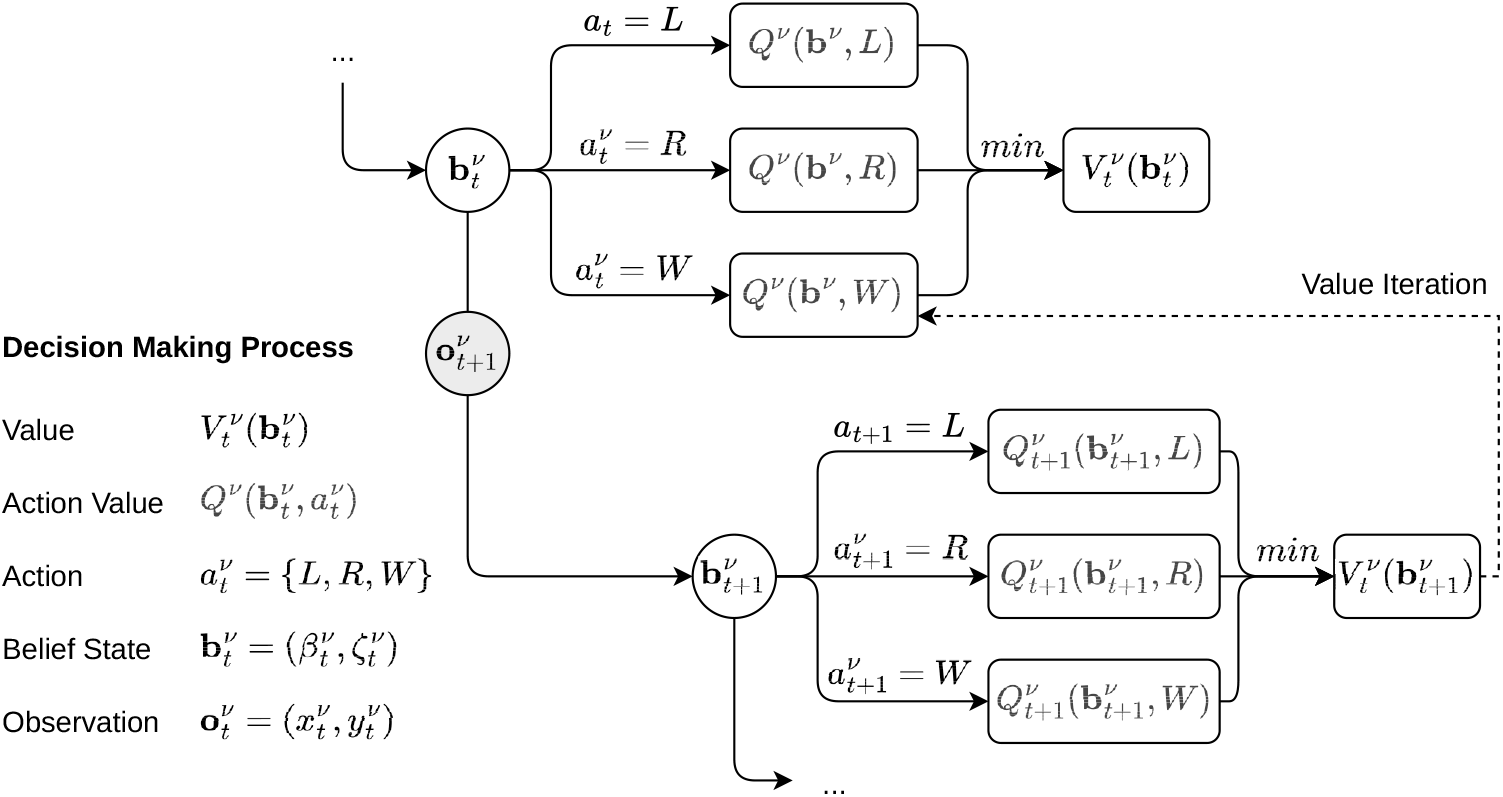
Optimal decision-making process in the POMDP model. At each time step *t* of trial *ν*, the belief state 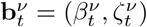 is updated using incoming go and stop observations 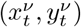. Action-values 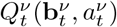 represent the expected long-run cost of each action. They are computed from the current belief state and intrinsic costs. The agent selects the action from Go-Left (*L*), Go-Right (*R*), and Wait (*W*) that minimizes this expected cost. The value of the Wait action is derived recursively via value iteration (dotted line). This applies the Bellman equation to back-propagate expected future costs from 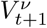 to 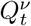.

If the participant chooses a terminal response Go-Left (*L*) or Go-Right (*R*) at time *t ≤ T*, the immediate cost incorporates three components: the accumulated time (*c*_*t*_ *· t*), the expected directional error on go trials (*c*_ge_), and the expected stop error on stop trials (*c*_se_):

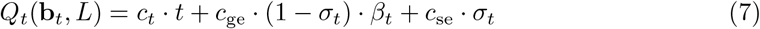

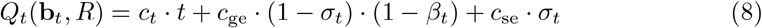

where the stop trial probability *σ*_*t*_ is derived directly from *ζ*_*t*_.

Alternatively, the participant can choose to Wait (*W*) to accumulate more information. If *W* is chosen consistently until the terminal step *T*, a timeout occurs. This incurs time costs and a go missing penalty (*c*_gm_):

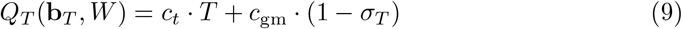

For any *t < T*, the value of waiting is derived via the Bellman optimality equation [33]. It is calculated from the expected optimal value of the next belief state, marginalized over all possible observations at the next timestep **o**_*t*+1_ = (*x*_*t*+1_, *y*_*t*+1_):

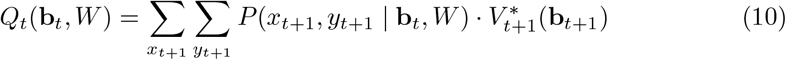

Here, 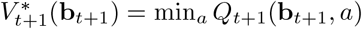 denotes the value of following the optimal policy from time *t* + 1 onward. The transition probabilities *P* (*x*_*t*+1_, *y*_*t*+1_ | **b**_*t*_, *W*) are computed by marginalizing over the future stop signal state *z*_*t*+1_ and the latent go direction *d*. The detailed expansion of these transition probabilities is provided in Section A.3 of S1 Appendix.

The complete set of action-values is computed recursively via backward induction from *T* down to *t* = 1. For computational implementation, the continuous belief space is discretized into an 80 × 80 grid using a tensor-based approach (see Section A.4 of S1 Appendix).

Finally, to capture the stochasticity of human behavior, we model the actual action selection using a softmax policy applied to the pre-computed optimal costs:

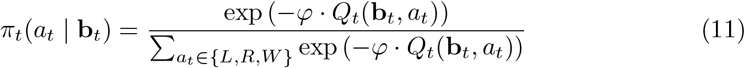

where the inverse temperature parameter *φ* controls the degree of determinism in the response. The negative sign ensures that lower expected costs generate higher choice probabilities.

### Estimation framework: TeSBI

Calculating the exact likelihood for the full POMDP model is computationally intractable due to the high-dimensional parameter space and the non-linear, discrete nature of the belief and action-value updates. Furthermore, observed behavioral sequences contain rich, multi-faceted temporal interactions (e.g., those induced by the SSD staircase) that are difficult to capture with simple summary statistics.

To address these challenges, we introduce Transformer-encoded Simulation-Based Inference (TeSBI). This approach extends standard SBI frameworks [25] by integrating a Transformer encoder [26] to learn compact, informative representations of the behavioral data. As illustrated in Fig 4, the TeSBI procedure consists of two stages: (1) Amortized training: mapping simulated behavioral sequences into embeddings via a Transformer encoder, and using a Neural Spline Flow (NSF) [24, 34] to map these embeddings to parameter densities; and (2) Posterior inference: sampling individual-level posterior parameters from observed behavioral sequences using the trained network.

**Fig 4.**
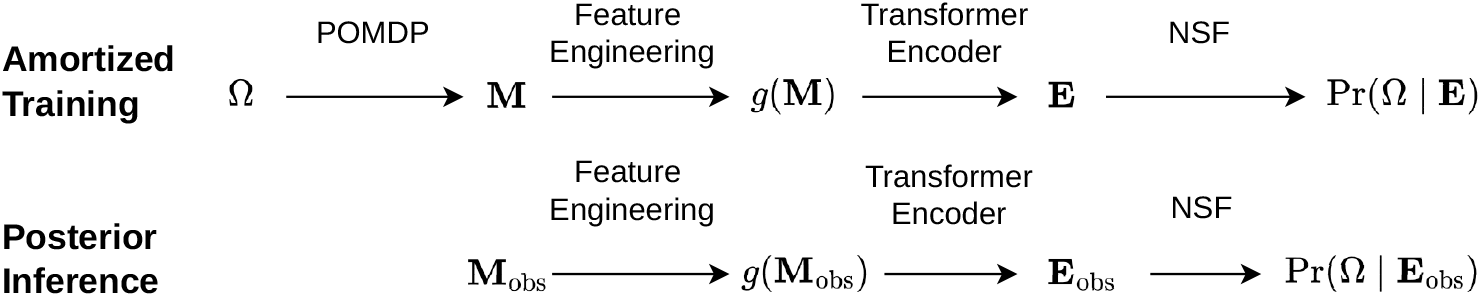
Overview of the end-to-end TeSBI procedure. (1) Amortized training: Parameters (Ω) sampled from the prior generate simulated behavior (**M**) via the POMDP simulator. After feature engineering, a Transformer encoder extracts temporal embeddings (**E**), and a Neural Spline Flow (NSF) is trained to map these embeddings to posterior densities. (2) Posterior inference: For each individual, the trained Transformer embeds their engineered observed sequence (**M**_obs_), and the NSF estimates the parameter posteriors.

We established model reliability through parameter recovery (verifying the identifiability of all parameters from synthetic data) and posterior predictive checks (PPCs, confirming that simulated behavior generated from inferred parameters matches empirical patterns). Detailed algorithmic procedures, network architectures, and hyperparameter configurations, and transformer encoded representations are provided in Section B of S1 Appendix.

### Amortized training

#### Feature engineering

To avoid the loss of information inherent to handcrafted summary statistics [35], we extract both trial-level and inter-trial features to prepare the full behavioral sequences as structured inputs for the downstream Transformer encoder. These arise from simulated sessions.

Each simulated session involves *N*_tr_ = 360 trials. For a given trial *ν*, the simulator produces a measurement tuple:

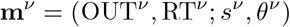

where *s*^*ν*^ is the trial type, *θ*^*ν*^ is the SSD on stop trials (*null* on go trials), and OUT^*ν*^ ***∈*** {GS, GE, GM, SS, SE} denotes the behavioral outcome categories (go success, go error, go missing, stop success, stop error). As for the adaptive staircase procedure used in the original ABCD study, the SSD increases or decreases by 2 time steps (i.e., 50 ms) following a stop success or stop error, respectively, and is restricted to the range of [2, 34] time steps (i.e., [50 ms, 850 ms]).

We denote the full session sequence as 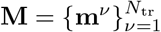. To capture temporal dynamics, we employ a feature engineering operator *g*(·) that augments each trial into an 18-dimensional vector:

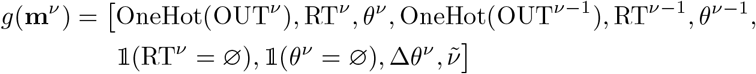

where OneHot(·) denotes a one-hot encoding of the 5 outcome types, and 1 (·) is an indicator function checking for missing values (∅). The vector also incorporates the previous trial’s features, the change in SSD step-size (Δ*θ*^*ν*^ = *θ*^*ν*^ *™ θ*^*ν™*1^), and a normalized trial index 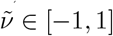 tracking the trial’s relative position within the session.

For the first trial, previous features are initialized with zeros. Applying this operator to the full session yields the finalized sequence 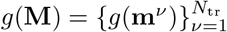,which is ready for downstream embedding.

#### Transformer encoder

To process these structured sequences, we employ a Transformer encoder *f*_*ϕ*_ as a feature extractor. Conceptually, we treat each trial as a *token* that contains information about the current behavior and context. The entire behavioral sequence acts as a *document*, the learned behavioral embedding serves as its *semantics*, and the underlying generative model parameters represent its core *themes*.

Formally, the encoder uses a form of self-supervision to map the transformed behavioral sequence into a dense summary embedding:

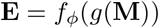

from which it is ultimately possible to recover the posterior over the parameters that generated the sequence. Through its multi-head attention mechanism, the encoder naturally captures complex temporal structures (e.g., SSD adaptation dynamics, post-error slowing, preparatory effects, and fatigue) without requiring manual specification. The dense embedding **E** then serves as the direct input for the downstream density estimator.

#### Neural Spline Flow

We approximate the posterior distribution *P* (Ω | **E**) using a Neural Spline Flow (NSF) density estimator *q*_*Ψ*_(Ω | **E**) [34]. Conditioned directly on the summary embedding **E** extracted by the Transformer, the NSF and the encoder are optimized jointly across a large offline dataset of pre-simulated behavior. By minimizing the negative log-likelihood of the ground-truth generative parameters given these embeddings, the unified network learns a global mapping from complex behavioral patterns to parameter probability densities. The density estimation pipeline was implemented using the zuko normalizing flow library [36].

### Posterior inference

As an amortized estimator, the fully trained TeSBI model performs zero-shot inference for all participants via a single forward pass. This eliminates the need for computationally expensive, subject-specific simulations.

For each participant *u*, the network directly maps their observed sequence 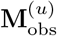 to a summary embedding 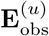. This embedding then conditions the NSF to estimate the individualized posterior:

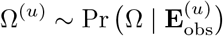

We use the posterior mean as the point estimate for subsequent analyses, computed via 1,000 samples per participant.

## Results

We characterize the behavior of the proposed POMDP framework and its application to the ABCD dataset. First, we illustrate the model’s theoretical properties. This includes the within-trial dynamics of perceptual inference, the derived optimal policy, and the resulting action-values. Next, we validate the reliability of the TeSBI fitting pipeline. Finally, we present the empirical results, focusing on the distribution of estimated parameters and their association with ADHD scores.

### Model simulations

To illustrate the internal dynamics of the proposed framework, we simulated behavior using our primary model (M6.1). Details regarding the selection of M6.1 (including model specification, parameter recovery, and posterior predictive checks) are provided in Section C of S1 Appendix.

#### Perceptual inference: belief states dynamics

The temporal evolution of the agent’s belief states reveals distinct patterns across five different trial outcomes (Fig 5). Here, we visualize example and summary simulations where the true go cue is Right and, on stop trials, the stop signal appears at SSD=11.

**Fig 5.**
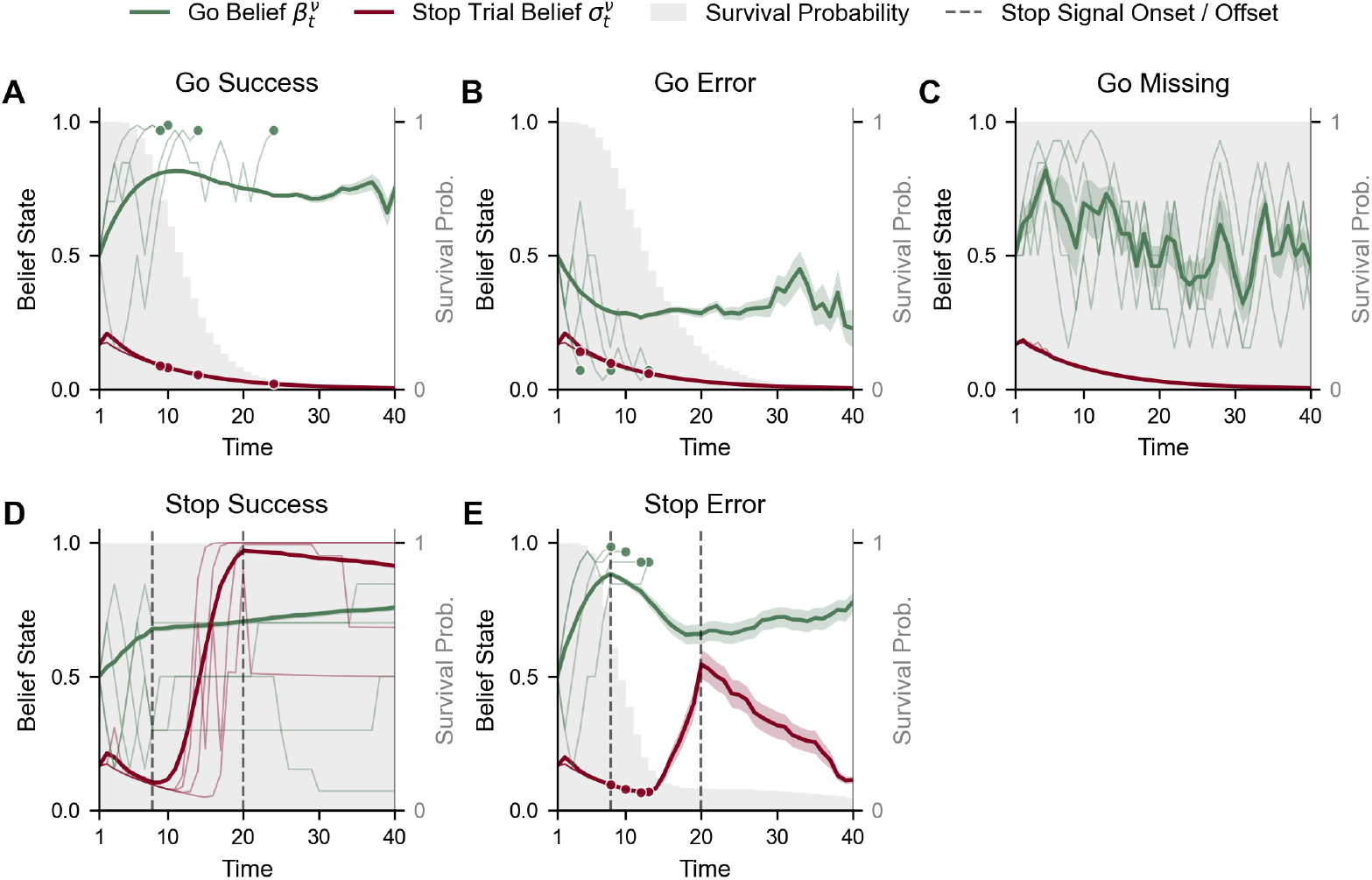
Belief state dynamics across five outcomes. Top row: go trials (with a right go cue). Bottom row: stop trials (with a right go cue and stop signal). Vertical dashed lines in the bottom panels indicate the stop signal window: onset at SSD=11 (200 ms) and offset after a duration of 12 time steps (300 ms). Thin lines depict four randomly sampled individual trajectories per outcome; solid dots mark the decision times (RTs). Thick curves represent the mean belief trajectory 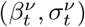 computed over ongoing trials at each time step (shaded bands: s.e.m.; 6, 000 go trials, 1, 200 stop trials). Note that mean curves may not reach the trial horizon because individual trajectories are truncated at the go-decision point. The gray shaded area depicts the survival probability, indicating the proportion of trials that have not yet ended. Parameters are from a representative participant (the 95th percentile goodness-of-fit) derived from our primary model (M6.1): 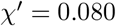, *χ* = 0.701, 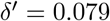 ***δ*** = 0.901, *c*_se_ = 4.705, *φ* = 44.653, *c*_ge_ = 1.647, *c*_gm_ = 15.782, *c*_*t*_ = 0.046, and 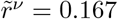.Although the fitted non-decision time is *τ* = 7 steps (175 ms) for this subject, it is set to 0 here to visualize the cognitive process starting from *t* = 1.

In Go Success trials, 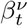 accumulates towards 1, reflecting successful direction recognition (the white area indicates the proportion of trials where the model had already chosen to go). Conversely, in Go Error trials, 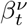 drifts below 0.5, indicating that the agent erroneously accumulated observations for the left direction despite the cue being right. In Go Missing trials, 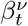 stagnates around 0.5, implying that the agent fails to resolve directional ambiguity. This prompts persistent waiting until the time limit. Across all go trials, the stop signal belief (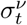, red line) consistently decays towards zero. In Stop Success trials (successful inhibition), 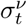 initially decays but surges sharply immediately after the stop signal onset (first vertical dashed line), accurately tracking the signal’s presence. It peaks before the signal offset and declines thereafter. In contrast, in Stop Error trials, the rise in 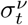 is notably delayed and attenuated.

Simultaneously, the go belief 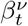 often rises rapidly, overpowering the weaker stop belief and leading to a premature go response before the stop signal is fully processed.

The dynamics of 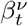 and 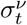 reflect how sensory evidence for the go cue and stop signal accumulates over time. This establishes the perceptual basis for the subsequent optimal control.

### Optimal control: probabilistic policy and action values

#### Probabilistic policy

Fig 6 illustrates the POMDP choice strategy and belief state evolution. A softmax function maps *Q*-values across the joint belief space go belief (*β*^*ν*^) and stop signal belief (*ζ*^*ν*^) into regions favoring Wait (light gray), Go-Left (yellow), or Go-Right (blue). As the deadline (*t* = 40) approaches, the go regions expand inward, reflecting an increased urgency to act as the opportunity cost of a timeout outweighs the risk of an error. The full policy evolution is detailed in Section A.5 of S1 Appendix.

**Fig 6.**
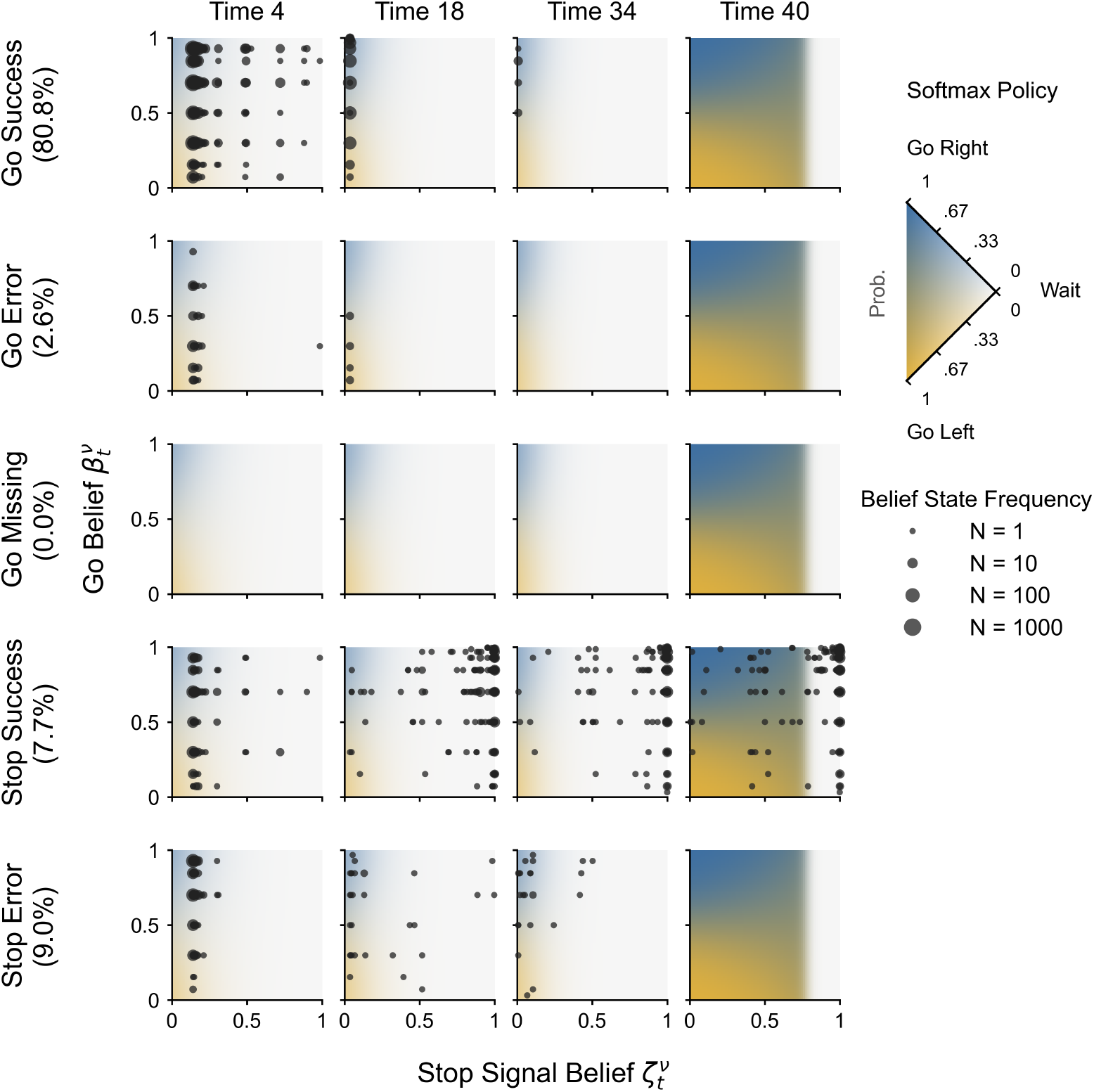
Softmax policy and belief state trajectories across selected time steps and trial outcomes. Background colors represent the softmax policy probabilities at selected time steps across the joint belief space: go belief 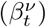 and stop signal belief 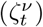. Because the softmax function applies a highly non-linear transformation that compresses extreme *Q*-values, the color mapping is visually enhanced using gamma correction (gamma is 0.05). The simplex legend on the right reflects this corrected mapping, indicating the probability distribution between Go-Right (blue), Go-Left (yellow), and Wait (light gray). Overlaid black dots represent the agent’s belief states, drawn from simulations of 6, 000 go trials (right go cue) and 1, 200 stop trials (right go cue; stop signal with SSD=11). The trials are categorized into five outcomes with their respective simulated frequencies. The size of the belief state dots represents their density. Trajectories terminate at the go-decision point, while states in Go Missing and Stop Success outcomes persist in the central Wait region until the deadline. Parameters are identical to those in Fig 5.

Realized actions depend on how belief states traverse this probabilistic space. In simulated trials (right-cue, SSD=11), go success and go error trajectories rapidly accumulate directional evidence along the *y*-axis, triggering early responses in the Go-Right or Go-Left regions, respectively. Rare go missing trials occur when weak evidence leaves the state stranded in the central Wait region.

On stop trials, successful inhibition (stop success) occurs when signal detection rapidly increases *ζ*^*ν*^, shifting the belief state rightward into the Wait region. Conversely, a stop error happens if go evidence accumulates too quickly, crossing the go threshold before the stop belief can sufficiently rise. Overall, these latent dynamics mutually corroborate the belief trajectories and action-values shown in Figs 5 and 7.

**Fig 7.**
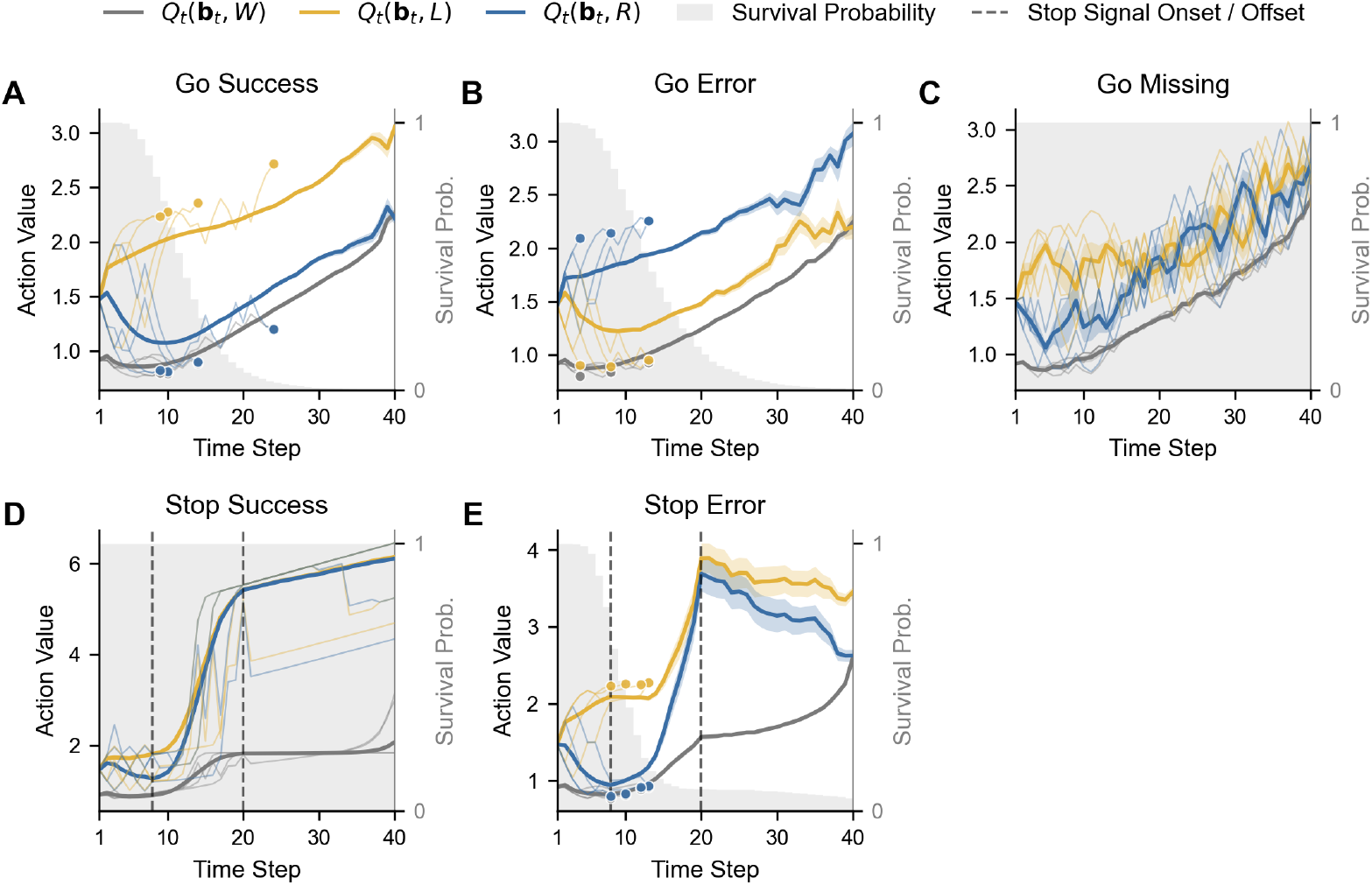
Action-value dynamics across five outcomes. Thick curves represent the mean action-value trajectory 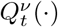 for Go-Left, Go-Right, and Wait actions, computed over ongoing trials at each time step (shaded bands: s.e.m.; 6, 000 go and 1, 200 stop trials). A softmax policy is used for decision-making: actions with lower expected costs (*Q*-values) are optimal and more likely to be chosen, though stochasticity permits the occasional selection of higher-cost actions. Plotting conventions (trial layouts, stop signal window, sample trajectories, decision markers, and survival probability) and all model parameters are identical to those in Fig 5.

#### Action-values

Action-values (*Q*-values) encode expected long-run costs. Under the stochastic softmax policy, lower *Q*-values correspond to higher selection probabilities. Simulated *Q*-values are shown in Fig 7, following the same conditions as in Fig 5.

In Go Success trials, the *Q*-value of Wait is initially dominant (i.e., lowest). As the directional belief becomes sufficiently certain, the *Q*-value of the correct action (e.g., Go-Right) decreases, narrowing the gap with that of Wait. This relative reduction rapidly shifts the softmax probability in favor of the action, triggering a Go-Right response. Importantly, because the policy is stochastic, actions can be triggered even before a strict crossover in values occurs, provided the probability mass has shifted sufficiently. Moreover, while this shift is sharp in individual simulations (thin lines), the averaged trajectories (thick curves) appear smoother due to temporal variability across trials.

In Go Error trials, the *Q*-value of the incorrect action (e.g., Go-Left) decreases erroneously due to noisy belief accumulation favoring the wrong direction, resulting in a directional error. In Go Missing trials, Wait consistently retains a significantly lower *Q*-value than the alternatives, keeping the probability of go actions negligible throughout the trial.

In Stop Success trials, Wait remains the optimal action (lowest *Q*-value) throughout the trial. Crucially, this sustained waiting reflects strategic inhibition under risk, not perceptual uncertainty. As shown in Fig 5, the agent quickly detects the stop signal (rapid rise in stop belief 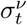); this high certainty, combined with the high cost of stop errors (*c*_se_), drives the *Q*-value of Wait down to enforce inaction (rather than prolonged uncertain waiting).

In Stop Error trials, the *Q*-value of a go action becomes competitive with Wait despite the stop signal. Multiple interacting factors, such as stronger early go directional observations, weaker stop signal detection, or a lower stop error penalty, collectively shift the action-value to favor execution over inhibition.

### Model selection and validation

To ensure parameter identifiability, we evaluate 15 candidate POMDP variants using an iterative step-down procedure. Starting from the unconstrained full model, we systematically reduce unidentifiable parameters by fixing those that fail the recovery test (Pearson’s *r <* 0.65) to specific posterior quantiles derived from the preceding model. We repeat this reduction process until all remaining free parameters are robustly identifiable.

We then evaluate the fit of these models using in-sample and out-of-sample posterior predictive checks (PPCs). We quantify the distance between behavioral observations and simulations via an aggregate distance metric comprising nine components: choice proportions (GS, GE, GM, SS), Wasserstein distances for RT distributions (GS, GE, SE), and Kolmogorov-Smirnov distances for RT distributions (GS, SE).

We select M6.1 as the primary model, as it yields the lowest total distance among the fully identifiable variants. It successfully recovers all six core parameters, showing strong correlations for go precision (*χ*, Pearson’s *r* = 0.89), non-decision time (*τ, r* = 0.83), time cost (*c*_*t*_, *r* = 0.82), and stop error cost (*c*_se_, *r* = 0.81), alongside reliable recovery for go error cost (*c*_ge_, *r* = 0.73), noting the rarity of go error behaviors; and stop precision (***δ***, *r* = 0.73), especially given the sparsity of stop trials in the task design.

To illustrate the predictive power of M6.1, Fig 8 shows fits for three participants representing Good, Moderate, and Poor fits (the 95th, 50th, and 5th percentiles of the aggregate distance, respectively). Across these performance levels, the model accurately reproduces the empirical outcomes, RT distributions (go and stop error), and task dynamics (SSD tracking and stop success rates). Even for the “Poor” fit participant, the model provides a reasonable approximation; the higher distance metric primarily reflects the participant’s inherently noisy behavior (e.g., non-monotonic inhibition across the SSD staircase) rather than structural model failures.

**Fig 8.**
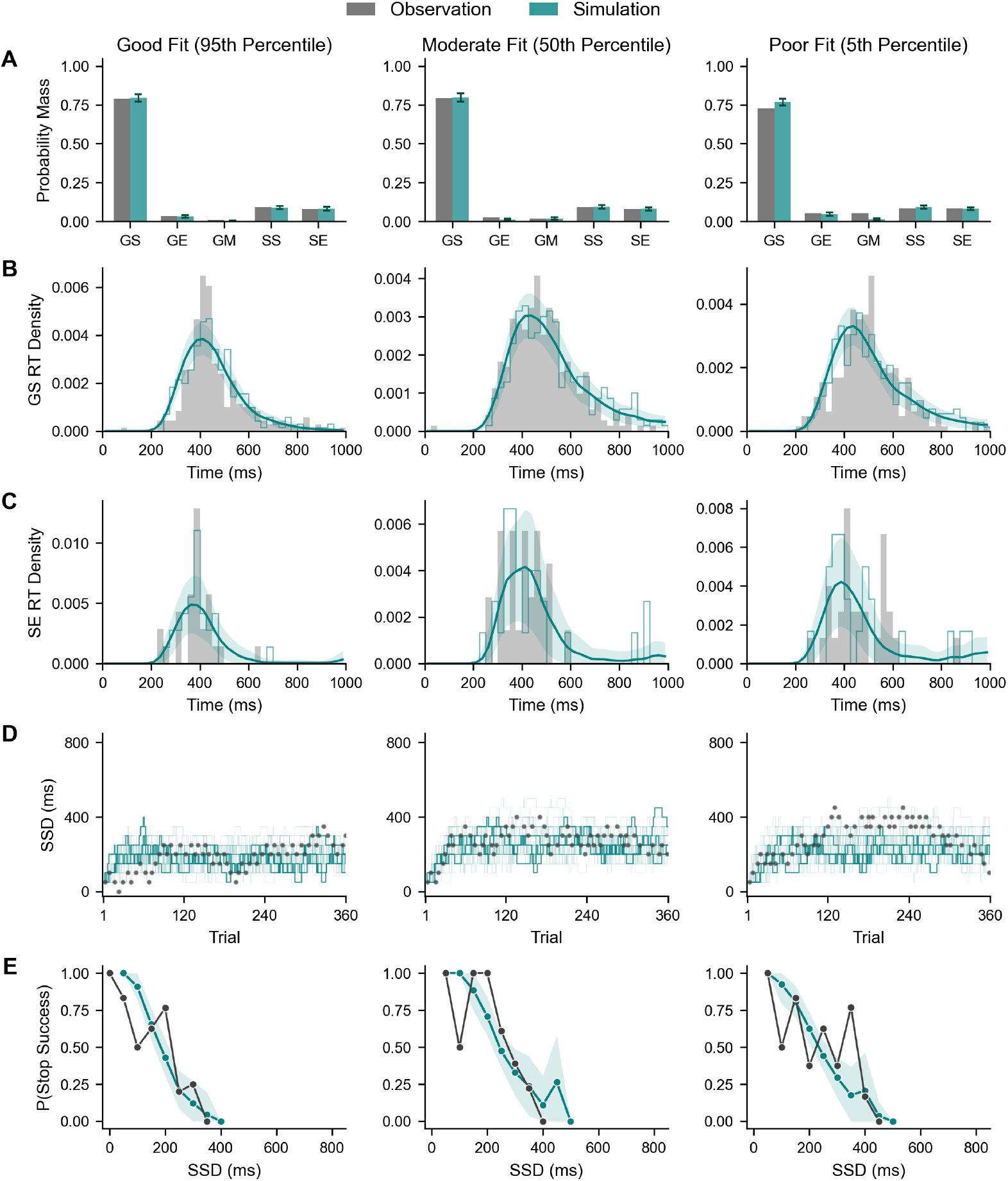
Representative posterior predictive checks for M6.1. Three participants illustrate Good (left), Moderate (middle), and Poor (right) model fits. (A) Observed outcome rates (gray) compared to the mean of 30 simulations (teal, generated using posterior mean parameters), with error bars indicating s.e.m. across simulations. (B–C) Observed reaction time distributions for Go Success and Stop Error (gray) compared to the mean simulated density (teal) and its variability (shaded area s.e.m.). A single simulation trace (thin teal line) is overlaid to illustrate a typical simulation. (D) SSD staircase dynamics. Observed trials (gray) are plotted alongside all simulated trajectories (faint teal), with three individual simulations (teal) highlighting the model’s adaptive behavior. (E) Inhibition dynamics. The observed probability of stop success (gray line and dots) is compared to the simulated mean (teal line and dots) and its variability (shaded area s.e.m.). All model time steps are converted to real time (1 time step = 25 ms).

Comprehensive details regarding the step-down model selection process, complete parameter recovery matrices, and PPCs results are provided in Section C of S1 Appendix.

Beyond individual parameter associations, we validate the holistic representations learned by the Transformer encoder using Canonical Correlation Analysis (CCA) and Principal Component Analysis (PCA). CCA reveals a strong, multidimensional correspondence between latent embeddings and model-agnostic behavioral metrics (primary variate *ρ*_*c*_ = 0.995, *p <* 0.001), confirming the encoding of both static and temporal task dynamics. Furthermore, PCA projections reveal that these behavioral fingerprints form a continuous computational manifold, confirming that higher ADHD scores are heterogeneously distributed across the population rather than clustered as an isolated deficit. Detailed cross-loadings and visualizations are provided in Section B.2 of S1 Appendix.

### Computational attributes and ADHD score associations

Here, we define the six core continuous parameters derived from our POMDP model collectively as latent computational attributes. These attributes quantify distinct aspects of an individual’s cognitive architecture, specifically capturing the characteristics of their perceptual processing and subjective valuation. To obtain these metrics, we draw 1, 000 samples from the posterior parameter distribution of each participant and compute the mean as an individual-level point estimate. The aggregate parameter distributions across the valid baseline year sample are provided in Section E.1 of S1 Appendix.

To investigate how computational attributes relate to clinical characteristics, we analyzed the association between the six estimated POMDP parameters and ADHD scores. We included sex, intelligence quotient (IQ, derived from the WISC Matrix Reasoning score [37]), and medication status (coded as 1 for stimulant prescription [38]) as established covariates.

To facilitate comparability, all continuous variables, including the estimated POMDP parameters, ADHD scores, and IQ scores, were standardized (*z*-scored) prior to the analysis. We then regressed each parameter on these predictors using multiple linear regression. To ensure model parsimony, we performed stepwise model selection via the Bayesian Information Criterion (BIC). Because BIC consistently penalized the inclusion of interaction terms across all parameters, we retained the simplified main-effects model for all subsequent analyses (see Section E.2 of S1 Appendix for detailed comparisons).

Within the main-effects model (Table 4), higher ADHD scores significantly predict lower go precision (*χ*; *β* = *™*0.068, *p <* .001), lower stop error cost (*c*_se_; *β* = *™*0.053, *p* = .004), lower time cost (*c*_*t*_; *β* = *™* 0.050, *p* = .007), and lower go error cost (*c*_ge_; *β* = *™* 0.043, *p* = .020). We observe no significant associations with stop precision (***δ***) or non-decision time (*τ*).

**Table 4.** Parameter Regression Coefficients and Effect Sizes.

| Parameters | Coef. | SE | p-value | Partial $R^2$ | $f^2$ |
| --- | --- | --- | --- | --- | --- |
| **ADHD** |  |  |  |  |  |
| $\chi$ | -0.0681 | 0.0182 | 0.0002*** | 0.0039 | 0.0039 |
| $\delta$ | -0.0038 | 0.0184 | 0.8351 | < 0.0001 | < 0.0001 |
| $c_{se}$ | -0.0533 | 0.0183 | 0.0036** | 0.0024 | 0.0024 |
| $c_{ge}$ | -0.0427 | 0.0184 | 0.0201* | 0.0015 | 0.0015 |
| $c_t$ | -0.0498 | 0.0183 | 0.0067** | 0.0021 | 0.0021 |
| $\tau$ | -0.0256 | 0.0184 | 0.1635 | 0.0005 | 0.0005 |
| **Sex (Male)** |  |  |  |  |  |
| $\chi$ | -0.0244 | 0.0336 | 0.4684 | 0.0001 | 0.0001 |
| $\delta$ | -0.0435 | 0.0340 | 0.2017 | 0.0005 | 0.0005 |
| $c_{se}$ | 0.2143 | 0.0338 | < 0.0001*** | 0.0111 | 0.0113 |
| $c_{ge}$ | -0.1094 | 0.0340 | 0.0013** | 0.0029 | 0.0029 |
| $c_t$ | 0.1584 | 0.0339 | < 0.0001*** | 0.0061 | 0.0061 |
| $\tau$ | 0.0006 | 0.0340 | 0.9866 | < 0.0001 | < 0.0001 |
| **IQ** |  |  |  |  |  |
| $\chi$ | 0.1455 | 0.0166 | < 0.0001*** | 0.0211 | 0.0216 |
| $\delta$ | -0.0374 | 0.0168 | 0.0264* | 0.0014 | 0.0014 |
| $c_{se}$ | 0.0362 | 0.0167 | 0.0306* | 0.0013 | 0.0013 |
| $c_{ge}$ | 0.0242 | 0.0168 | 0.1496 | 0.0006 | 0.0006 |
| $c_t$ | 0.0492 | 0.0168 | 0.0033** | 0.0024 | 0.0024 |
| $\tau$ | 0.0534 | 0.0168 | 0.0015** | 0.0028 | 0.0028 |
| **Medication** |  |  |  |  |  |
| $\chi$ | 0.2782 | 0.0789 | 0.0004*** | 0.0035 | 0.0035 |
| $\delta$ | -0.0793 | 0.0799 | 0.3210 | 0.0003 | 0.0003 |
| $c_{se}$ | 0.1109 | 0.0794 | 0.1628 | 0.0005 | 0.0005 |
| $c_{ge}$ | 0.1618 | 0.0798 | 0.0426* | 0.0012 | 0.0012 |
| $c_t$ | 0.0706 | 0.0796 | 0.3752 | 0.0002 | 0.0002 |
| $\tau$ | -0.0655 | 0.0798 | 0.4122 | 0.0002 | 0.0002 |

Note: $N = 3,566$ . \* $p < .05$ , \*\* $p < .01$ , \*\*\* $p < .001$ .

Covariates exhibit distinct computational profiles. Males show higher stop error (*c*_se_) and time costs (*c*_*t*_), alongside lower go error costs (*c*_ge_) compared to females, with no significant differences in precision or non-decision time. Higher IQ robustly predicts higher go precision (*χ*) and longer non-decision time (*τ*), along with higher stop error and time costs, though it is associated with slightly lower stop precision (***δ***). Medication status shows significant positive associations with go precision and go error cost.

Notably, while statistically robust, absolute effect sizes remain generally small (partial *R*^2^ *<* 0.015, Cohen’s *f* ^2^ *<* 0.015). However, ADHD characteristics and established covariates (Sex and IQ) exhibit comparable, parameter-specific effect size profiles.

Beyond mechanistic insights, POMDP parameters explain slightly more variance in ADHD scores than classic behavioral metrics after controlling for covariates. Adding these parameters to classic metrics significantly improves overall model fit, demonstrating they capture unique clinical information missed by conventional summaries (see Section E.3 of S1 Appendix).

We can illustrate these relationships by visualizing the distributions of the computational attributes stratified by ADHD scores and sex (Fig 9). As expected from the regressions, we observe population-level shifts across the clinical scores. For instance, the parameters significantly associated with ADHD, go precision (*χ*), stop error cost (*c*_se_), go error cost (*c*_ge_), and time cost (*c*_*t*_), highlighted with dashed mean lines, exhibit leftward shifts as ADHD scores increase. The distributions also highlight robust sex differences in specific cost valuations. Across all ADHD severity bins, males consistently show higher *c*_se_ and *c*_*t*_, and lower *c*_ge_ compared to females, maintaining a visible gap between their respective group means. However, although these shifts are highly statistically significant given the large sample size, they appear visually subtle due to the small absolute effect sizes.

**Fig 9.**
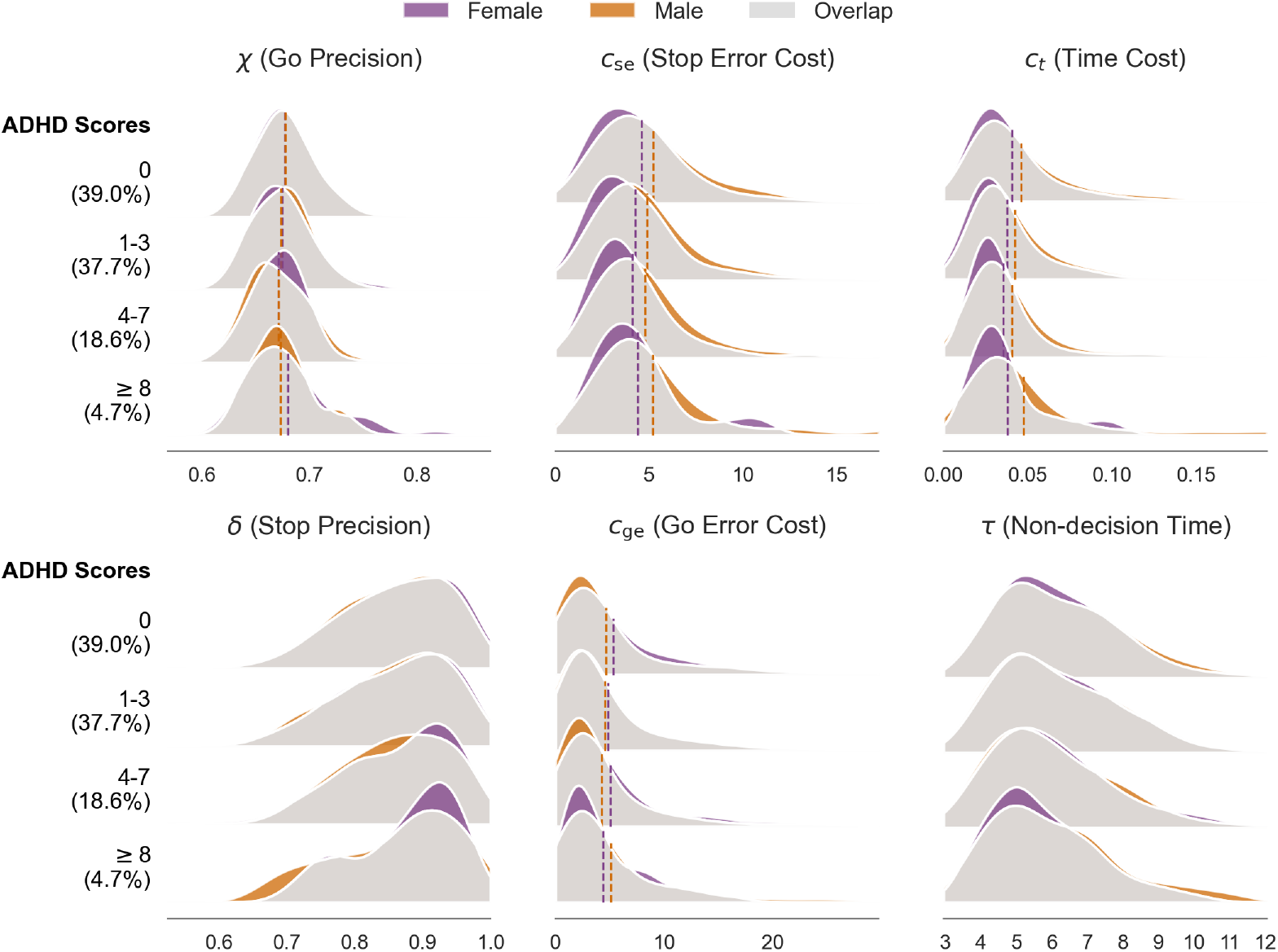
Distributions of participant-level computational attributes by ADHD scores and sex. Posterior distributions of the six POMDP parameters across the valid baseline sample (*N* = 3, 567). Participants are categorized into four clinical bins based on their ADHD scores (0, 1-3, 4-7, *≥* 8). Density plots are colored by sex, with the overlapping intersection shaded in gray. Vertical dashed lines denote the group-specific means for the four parameters exhibiting significant main effects of ADHD. The visualization highlights three key features of the computational attributes: progressive mean shifts across the clinical gradient, with four parameters showing significant decreases as ADHD scores increase (go precision *χ, p <* .001; *c*_se_ and *c*_*t*_, *p <* .01; *c*_ge_, *p <* .05); robust sex differences confined to the three cost valuations (*c*_se_, *c*_ge_, *c*_*t*_, all *p <* .01); and general individual heterogeneity, evidenced by wide probability densities and extensive overlap even between extreme clinical subgroups.

Crucially, the plots reveal substantial within-group heterogeneity. The wide probability densities and the extensive overlap between distinct clinical and demographic subgroups emphasize that ADHD is computationally heterogeneous. Rather than forming a homogeneous cluster, individuals with identical clinical scores can possess vastly different underlying cognitive architectures.

In summary, the computational attributes of higher ADHD scores stem from degraded information processing and altered cost valuation. This involves a reduction in go cue directional precision, alongside a blunted sensitivity to both errors and time.

Furthermore, the systematic exclusion of interaction terms demonstrates that these core computational alterations remain highly consistent regardless of patient sex, IQ, or medication status.

## Discussion

### Summary

We develop a Partially Observable Markov Decision Process (POMDP) model of the Stop Signal Task (SST) and apply it to the large-scale ABCD baseline dataset.

Following previous work formalizing inhibitory control as a continuous process of Bayesian inference and value-based action [21], our model unifies perception and decision-making within a single framework. To overcome the computational bottleneck of fitting such complex models, we introduce Transformer-encoded Simulation-Based Inference (TeSBI), an end-to-end pipeline enabling efficient and reliable parameter estimation at scale.

Applying this pipeline to the ABCD cohort, we systematically reduce the number of free parameters to ensure identifiability via parameter recovery and posterior predictive checks, resulting in a robust six-parameter POMDP model. We obtain subject-level parameters as computational attributes, which are then regressed on ADHD scores while controlling for sex, IQ, and medication status. Our main-effects regression reveals that, on average, higher ADHD scores are most robustly associated with a reduction in go cue directional precision, alongside a blunted sensitivity to stop error, go error and time costs.

Crucially, while these regressions capture overarching group-level trends, our latent behavioral embeddings reveal that these computational attributes form a continuous spectrum rather than discrete clinical clusters. The broad dispersion of individuals with higher ADHD scores across this space highlights substantial heterogeneity in their cognitive strategies, nuances that simple group-level averages inherently overlook.

### Mechanistic insights

The dominant accounts of the Stop Signal Task, initiated by the independent race model [4, 5] and its interactive variants [13, 14, 39], have provided profound insights into the architecture of cognitive control. By conceptualizing inhibition as a mechanical race between Go and Stop processes, these frameworks established the foundational methodology of using summary statistics to estimate the unobservable Stop Signal Reaction Time (SSRT) [40].

Building upon this foundation, advanced parametric extensions have further refined our understanding. Models such as the Bayesian estimation of Ex-Gaussian SSRT distributions (BEESTS) [41, 42], the racing diffusion Ex-Gaussian (RDEX-ABCD) model [43], the racing diffusion model (RDM) [44, 45] (which integrates the drift diffusion model (DDM) [7, 12, 46, 47]), and the linear ballistic accumulator (LBA) for the SST [48, 49], offer greater precision by fitting the full distributions of reaction times and accuracy. These approaches demonstrate how the *accumulate-to-bound* framework can capture the temporal dynamics of finishing times. However, by focusing primarily on perceptual accumulation, they typically abstract away the subjective valuation process, where an agent continuously weighs the cost of delaying an action against the risk of making an error at each moment.

Our POMDP framework explicitly addresses this gap by extending the foundational conceptualization of Shenoy and Yu [22], formalizing how the well-documented *race-like* dynamics of inhibition can naturally emerge from dynamic belief updates and continuous value optimization. This conceptual shift provides a normative perspective on how subjective sensory uncertainty and intrinsic costs optimally shape these racing processes. This approach offers three key advantages.

First, regarding *perceptual inference*, we augment the original framework by introducing subjective sensory noise to explicitly capture perceptual ambiguity.

Furthermore, we generalize the model structure to accommodate a dependent SST design, in which the stop signal masks the go cue. By modeling this interaction such that the agent’s rising belief in the stop signal mathematically degrades its certainty regarding the go cue, the perceptual masking effect emerges as a natural property of Bayesian inference. Ultimately, this dependent framework subsumes the traditional independent processing assumption as a special case.

Second, regarding the *optimal policy*, our model relies on dynamic, value-based action valuation rather than fixed decision thresholds. While standard models such as the DDM constitute the asymptotically optimal policy for simple 2AFC tasks [19, 50, 51], the continuous uncertainty and conflicting goals of the SST require the moment-by-moment, value-driven trade-off between the different possible trial types to be captured. From this perspective, reaction time slowing is interpreted as an active, strategic adjustment to minimize expected costs, rather than a passive braking mechanism. Section D of the S1 Appendix provides a qualitative comparison between our value-based mechanism and a more standard accumulate-to-bound architecture (RDEX-ABCD), a race model adapted to the peculiarities of the ABCD task [43]. The models fit comparably well, opening an important question for the future in mapping their rather different parameterizations onto each other.

Crucially, our computational perspective offers new clinical insights. While prior studies have highlighted stop-signal perceptual deficits as a plausible mechanism underlying ADHD [9], our findings tentatively suggest an alternative perspective (albeit with very small effect sizes). Within a unified framework like the POMDP, we hypothesize that a substantial portion of the behavioral variance traditionally attributed to pure perceptual impairments might also stem from top-down alterations in how individuals subjectively weigh the costs of errors and delays. Similarly, while recent DDM-based analyses of the ABCD task link ADHD to longer non-decision times [43], we observed only a non-significant negative trend. This divergence likely arises because our included covariates (sex, IQ, and medication) captured significant variance in non-decision time, combined with the limited clinical ADHD severity in the ABCD study.

Finally, regarding computational tractability, the introduction of *TeSBI* addresses the computational challenges that have typically restricted the application of complex cognitive models to large-scale cohorts. Unlike conventional summary-statistic methods, our Transformer-based encoder captures the sequential dependencies of trial-by-trial data. This enables the efficient and reliable estimation of individual-level posterior distributions, facilitating high-throughput computational phenotyping.

### Limitations and future directions

Our study has several limitations that highlight important avenues for future research. These can be broadly categorized into modeling and inference constraints, data characteristics, and clinical applications.

First, regarding modeling constraints, we assume deterministic time perception (i.e., known trial duration *T*). In reality, however, human time perception is inherently noisy [52]. Incorporating stochastic timing 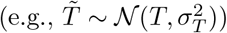 would allow agents to act based on an approximation of the remaining time. This adjustment would smooth the value transitions near the deadline and better capture the variability of premature and late responses.

Second, our current generative model omits the dynamic anticipation of stop signals. Within a trial, we do not model the participant’s prior expectation of exactly when the stop signal would appear (assuming a constant within-trial hazard rate *λ*). Across trials, we also abstract away how sequential history influences prior beliefs. While the SSD is primarily determined by a self-adaptive staircase procedure based on previous Stop outcomes, the number of consecutive Go trials also indirectly shapes behavior.

Specifically, a long sequence of Go trials may initially decrease the predicted probability of an upcoming Stop trial (via Bayesian updating [20, 22, 53, 54]) and subsequently increase it as participants expect Stop trials to recur. However, because the 1/6 stop ratio in the ABCD task restricts the fluctuation of this prior, fixing it to 1/6 allows us to focus on within-trial dynamics. As shown in our sensitivity analysis (Section A.6 of S1 Appendix), variations within this restricted range generate small but observable shifts in performance; thus, fixing the prior simplifies the current model. This approach establishes stable within-trial parameters first, allowing future extensions to free the stop prior and investigate across-trial effects.

Third, concerning inference constraints, we generate all simulated behaviors upfront before model training. Although we utilize parallel computing, matrix vectorization, and early stopping to accelerate the process, simulating data on CPUs remains a bottleneck compared to gradient-based training on GPUs. Future implementations could adopt an active learning or multi-round approach (i.e., iteratively generating data, training the model, and evaluating results) to further optimize computational resources, especially for models that feature dynamic learning across trials.

Regarding clinical relevance, we rely on ADHD scores assessed via the CBCL questionnaire rather than formal clinical diagnoses. The ABCD cohort is generally healthy, with relatively few participants exhibiting severe ADHD symptoms (see Section F of S1 Appendix for statistics). While this approximates the distribution in the general population, the lack of severe cases might lead to an underestimation of ADHD-related effects. Validating these computational attributes in clinical datasets with a larger proportion of diagnosed individuals is necessary to confirm the robustness of our results.

Furthermore, our current analysis is cross-sectional, relying exclusively on baseline data. Because the ABCD study features a longitudinal design, future work can track the developmental trajectories of these computational attributes and ADHD effects over a longer period (e.g., the ten-year span of the ABCD protocol). This offers a unique opportunity to investigate whether specific model parameters can serve as early predictors of symptom progression or as treatment biomarkers [55].

Finally, while this study focuses strictly on behavior, the availability of comprehensive neuroimaging data in the ABCD study is a tempting opportunity. Future research can link these computational parameters to specific neural correlates, potentially establishing them as quantitative endophenotypes that bridge the gap between underlying neural dysregulation and clinical behavior [56].

## Conclusion

In total, we built a POMDP model able to accommodate the rather particular version of the stop signal task implemented in the ABCD study, alongside an inference pipeline capable of fitting participants at an appropriate scale. Along with illustrating some of the peculiarities of this version, we showed significant but subtle correlations between the fit parameters and the rather limited range of ADHD scores in the study.

## Supporting information

### S1 Appendix. Supplementary methods and results

This appendix provides comprehensive details supporting the main contents, including: (A) Mathematical details of the POMDP model, covering derivations of stop process inference, the formulation for independent stop signal tasks, transition probabilities, tensor-based value iteration, full probabilistic policy visualization, and stop prior sensitivity analysis;(B)TeSBI architecture and validation, detailing the base architecture and transformer-encoded representations; (C) POMDP model selection and validation, including model specification, model selection procedure, parameter recovery analysis, and posterior predictive checks; (D) Comparison with alternative models; (E) Computational attributes, including aggregate distributions, parameter regression and interaction analysis, and predictor sets comparison; and (F) Model-agnostic statistical analysis.

## Acknowledgments

We thank Prof. Jakob Macke and Dr. Cornelius Schröder for their valuable advice on simulation-based inference. We also thank our colleagues from both laboratories for their support and the wider research community for inspiring discussions at conferences. We acknowledge the use of Gemini (Google) for assistance with LaTeX formatting and code correction.

## S1 Appendix: Supplementary Methods and Results

## A Mathematical details of the POMDP model

### A.1 Derivation of stop process inference

This section details the recursive Bayesian updates for the intermediate stop-signal belief 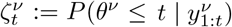 and the stop-trial belief 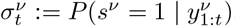.

1. **Adjusted hazard rate** 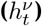 First, we derive the adjusted hazard rate, which represents the probability that the stop signal occurs at time *t*, given it has not occurred previously. This term must account for the uncertainty regarding the trial type *s*^*ν*^.

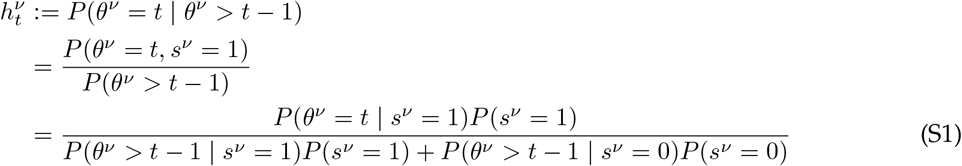

Using the geometric prior *P* (*θ*^*ν*^ = *t*|*s*^*ν*^ = 1) = *λ*(1 *™ λ*)^*t™*1^ and noting that for go trials (*s*^*ν*^ = 0), the stop signal never occurs (equivalent to *θ*^*ν*^ *→ ∞*, thus *P* (*θ*^*ν*^ *> t ™* 1|*s*^*ν*^ = 0) = 1), we obtain:

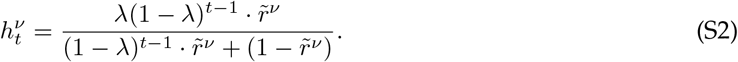
2. **Stop signal belief prediction** 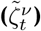 Before observing 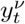, the agent computes the *predicted* belief that the signal has occurred by time *t*. This combines the previous posterior with the probability of a new onset:

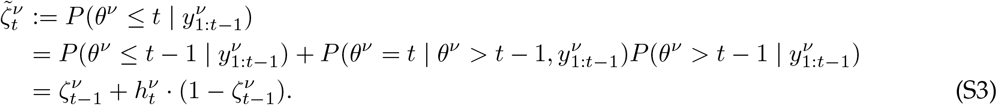

Complementarily, the probability that the signal has *not* occurred by time *t* is:

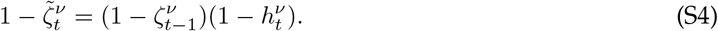
3. **Stop signal belief correction** 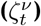Upon observing 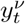, we update the belief via Bayes’ rule.

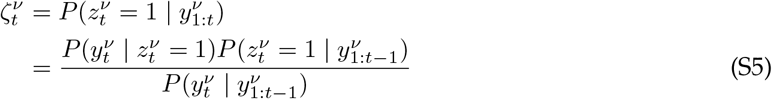

Expanding the denominator using the law of total probability and substituting Eq. S4:

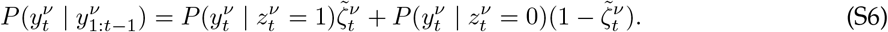

This yields the update equation used in the main text.
4. **top trial belief** 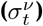Finally, the probability that the current trial is a stop trial is the sum of two disjoint possibilities: (a) the signal has already occurred, or (b) the signal has not occurred yet but the trial is essentially a stop trial (signal will occur in the future).

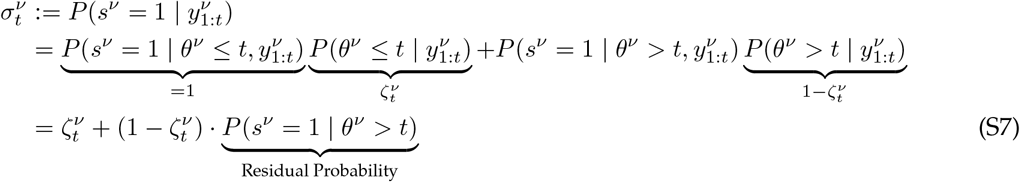

The residual probability represents the likelihood that a trial is a stop trial given no signal has appeared by time *t*:

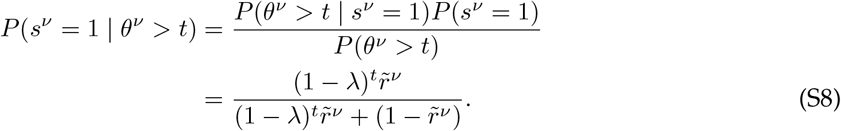

Combining these terms yields the final expression for 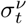.

### A.2 Formulation for independent stop signal task

In standard SST designs (e.g., auditory stop signals), the stop signal does not visually mask or interrupt the go cue. Consequently, the two sensory streams are effectively independent. Our framework accommodates this by removing the conditional dependency of the go observation on the stop signal state.

#### 1. Observation function for go process

Unlike the main model where the stop signal 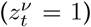 alters the go observation process, here we assume 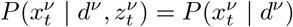. The go cue remains visible and unmasked throughout the trial. Hence, the observation probabilities are time-invariant and independent of stop signal onset (Table A).

**Table A.** Observation likelihoods for go process (independent SST)

| State $d^\nu$ | $P(x_t^\nu = L \mid d^\nu)$ | $P(x_t^\nu = R \mid d^\nu)$ | $P(x_t^\nu = N \mid d^\nu)$ |
| --- | --- | --- | --- |
| 0 | $(1 - \chi')\chi$ | $(1 - \chi')(1 - \chi)$ | $\chi'$ |
| 1 | $(1 - \chi')(1 - \chi)$ | $(1 - \chi')\chi$ | $\chi'$ |

#### 2. Decoupled inference and transitions

Under this independence assumption, the belief update for the go process no longer requires marginalization over the stop signal belief *ζ*^*ν*^. The update simplifies to:

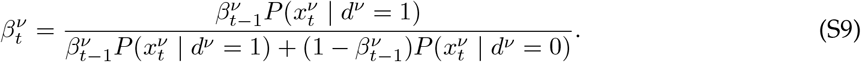

Consequently, the joint transition probability factorizes into two independent components:

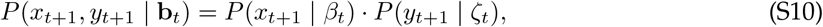

where *P* (*x*_*t*+1_ | *β*_*t*_) depends solely on the go belief, and *P* (*y*_*t*+1_ | *ζ*_*t*_) depends solely on the stop process dynamics. This formulation recovers the standard assumption of parallel, independent processing channels used in race models.

### A.3 Transition probabilities

To evaluate the action-value of Wait (*W*), the model must predict future observations by marginalizing over latent states. The transition probabilities *P* (*x*_*t*+1_, *y*_*t*+1_ | **b**_*t*_, *W*) are derived by marginalizing over the future stop signal state *z*_*t*+1_:

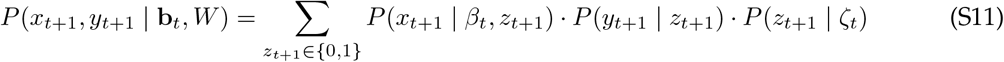

where the transition probability for the go observation *x*_*t*+1_ marginalizes over the latent true direction *d*:

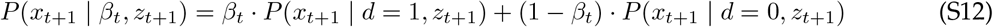

Given the approximation that the stop signal is functionally persistent once it appears, the transition probabilities for the stop signal state are given by the predicted belief 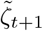.

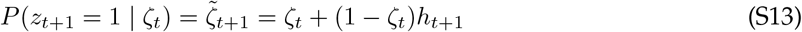

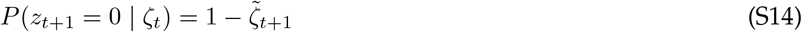

The primitive observation likelihoods *P* (*x*| *d, z*) and *P* (*y*| *z*) correspond to the functions defined in Tables 2 and 3 of the main text.

### A.4 Tensor-based value iteration

To enable scalable inference over the ABCD cohort, we vectorized the POMDP value iteration using tensor operations over a discretized belief space. This replaces slow, sequential loop-based computations with highly optimized matrix operations.

#### 1 Discretized belief grid

1. We discretize the continuous belief state space (*β, ζ*) *∈* [0, 1]^2^ into a uniform grid of resolution *N*_*β*_ *× N*_*ζ*_ (in our implementation, *N*_*β*_ = *N*_*ζ*_ = 80).

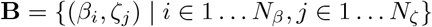

At any time step *t*, the value function *V*_*t*_ and action-value functions *Q*_*t*_(*·, a*) are represented as matrices of shape *N*_*β*_ *× N*_*ζ*_.

#### 2. Transition tensor construction

The core computational bottleneck is the expectation over future observations. We precompute a transition tensor *T*_*t*_ that encodes the probability of transitioning to any future observation pair (*x*^*′*^, *y*^*′*^) given any current belief state on the grid:

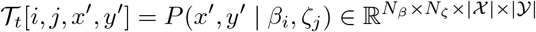

where |X |= 3 (Left, Right, Null) and |Y|= 3 (Absence, Presence, Null). This tensor is computed via the factorized likelihoods described in Eq. S11, efficiently broadcasting the independent updates for the Go process (dependent on *β*) and Stop process (dependent on *ζ*).

#### 3. Vectorized Bellman update

The expected value of waiting, *Q*_*t*_(**b**, *W*), involves an integral over future beliefs. In the discretized space, this becomes a tensor contraction. Let (*β*_*i*_, *ζ*_*j*_, *x*^*′*^, *y*^*′*^) be the deterministic next-step belief indices given current state (*i, j*) and observation (*x*^*′*^, *y*^*′*^). We compute the next-step value tensor of shape *N*_*β*_ *× N*_*ζ*_ *×* |*X* | *×* |*Y*| by gathering values from *V*_*t*+1_ according to the updated belief indices. The Q-value tensor for the Wait action is then computed in a single operation:

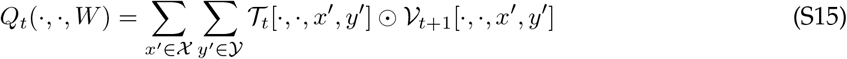

where ⊙ denotes element-wise multiplication. This vectorization allows us to solve the full dynamic programming problem for a single participant in milliseconds.

### A.5 Full probabilistic policy visualization

This section provides a comprehensive overview of the probabilistic policy and belief state evolution across all 40 time steps, categorized by the five possible trial outcomes.

The background heatmaps illustrate the softmax policy across the entire joint belief space, comprising the go belief 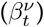 and stop signal belief 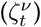, mapping regions that favor Wait (light gray), Go-Left (yellow), or Go-Right (blue). Heatmap colors are gamma-corrected for visual clarity (see Optimal control in the Results section for details). As the deadline approaches, the go regions progressively expand inward, reflecting the model’s increasing urgency to act to avoid a timeout.

Overlaid scatter points trace the simulated belief trajectories from 6, 000 go trials (right go cue) and 1, 200 stop trials (right go cue, SSD=11), with dot size representing the density of belief states. Categorizing these trajectories by outcome reveals the distinct patterns of observation accumulation driving the model’s behavior. Rapid vertical shifts lead to early responses in Go Success and Go Error trials; weak directional observations leave states stranded in the Wait region during Go Missing trials; and rapid rightward shifts upon Stop signal detection successfully suppress responses in Stop Success trials, provided the initial Go observations did not accumulate fast enough to cause a Stop Error.

**Fig A.**
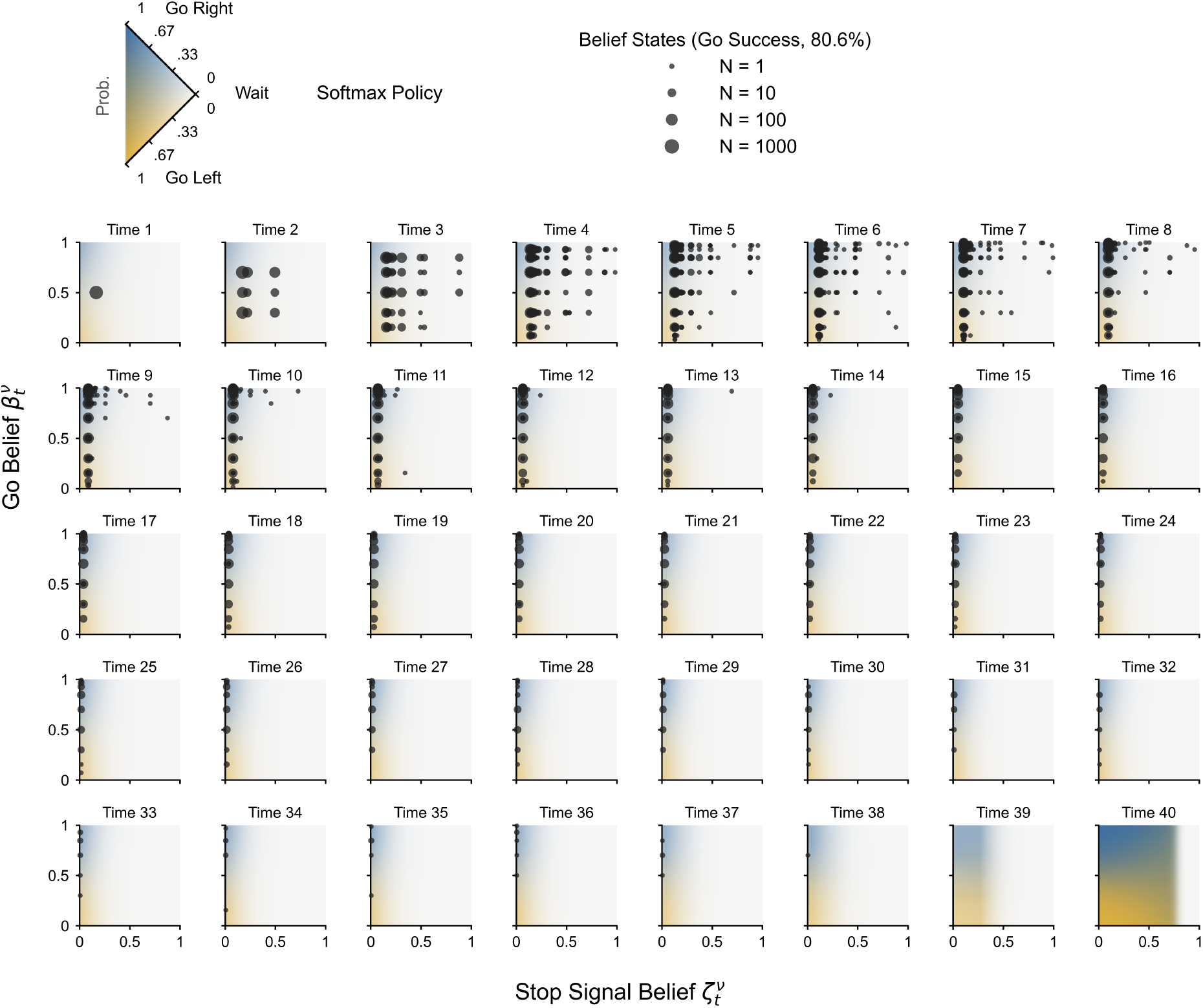
Softmax policy and belief states for Go Success trials. Background heatmaps represent action probabilities from the softmax policy. Scatter points show simulated trajectories from 6, 000 go trials (right go cue).

**Fig B.**
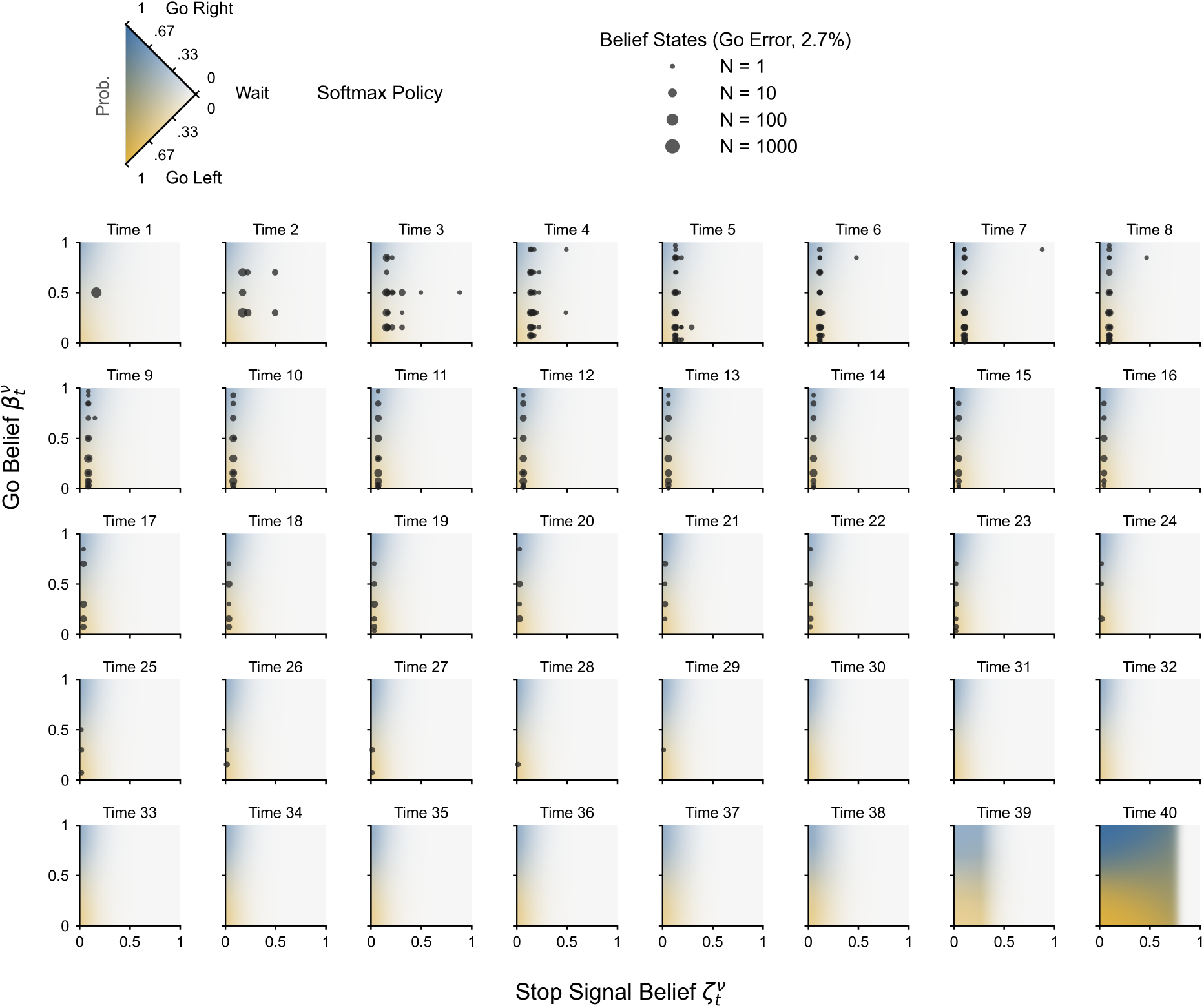
Softmax policy and belief states for Go Error trials. Background heatmaps represent action probabilities from the softmax policy. Scatter points show simulated trajectories from 6, 000 go trials (right go cue).

**Fig C.**
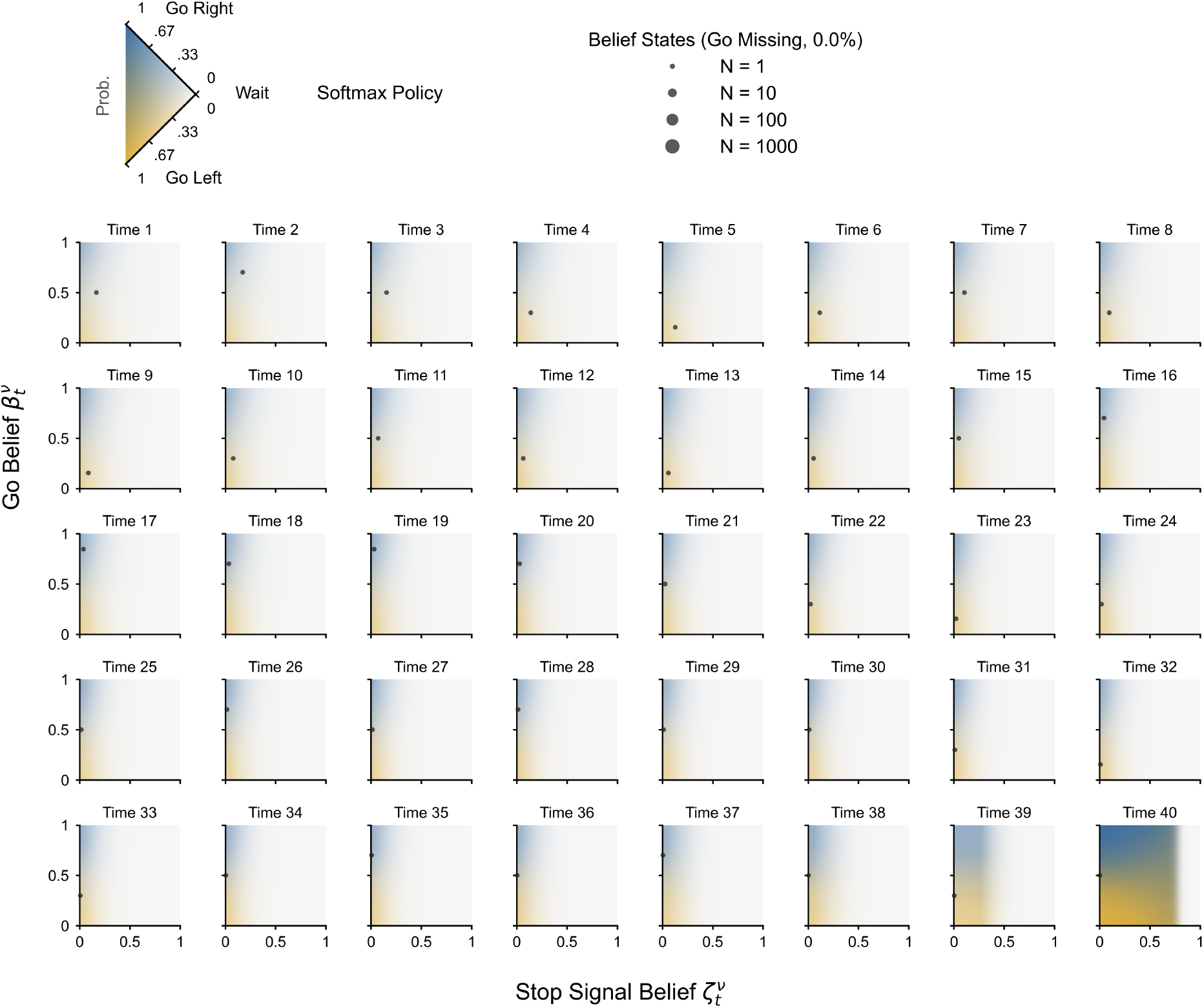
Softmax policy and belief states for Go Missing trials. Background heatmaps represent action probabilities from the softmax policy. Scatter points show simulated trajectories from 6, 000 go trials (right go cue).

**Fig D.**
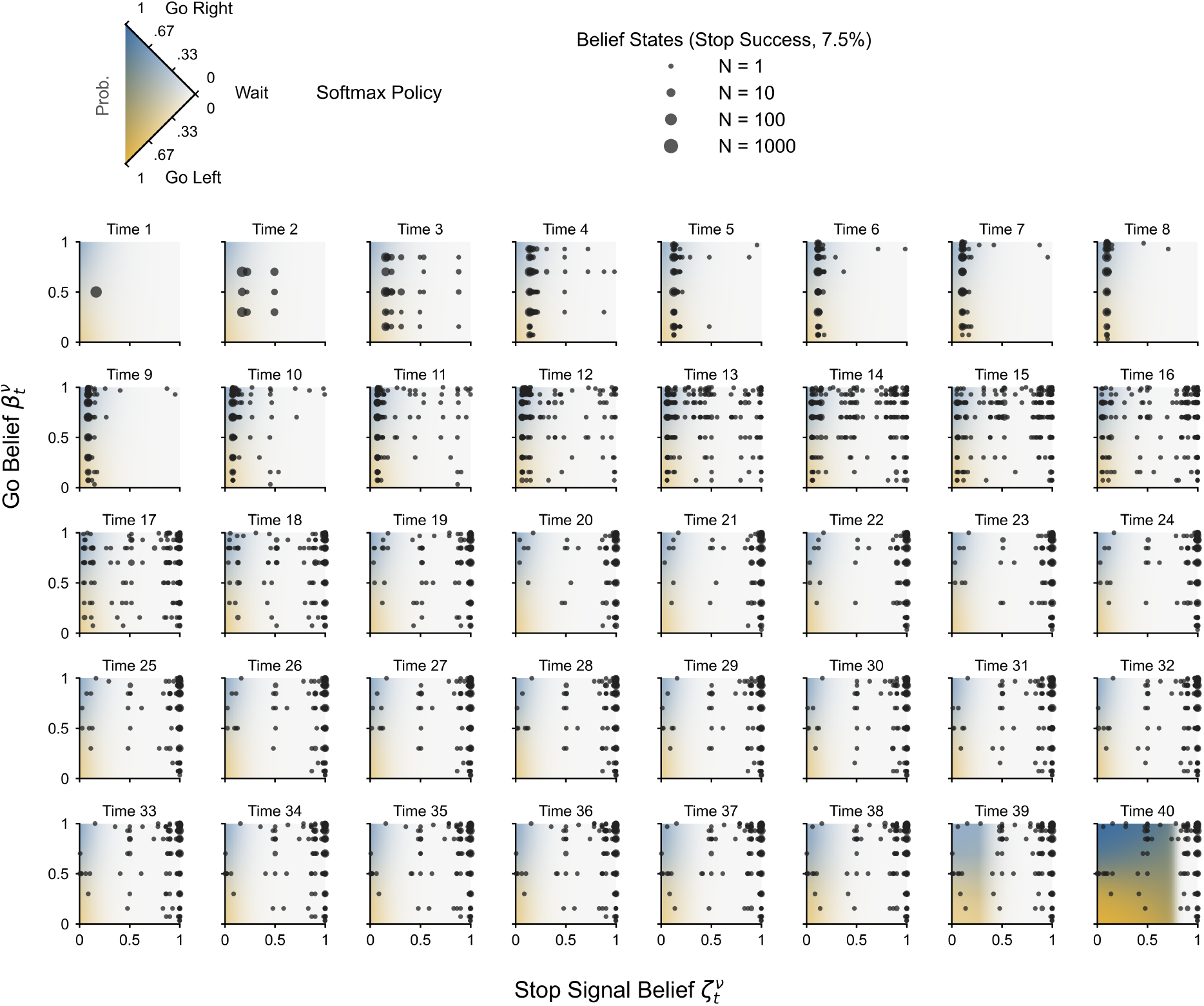
Softmax policy and belief states for Stop Success trials. Background heatmaps represent action probabilities from the softmax policy. Scatter points show simulated trajectories from 1, 200 stop trials (right go cue, SSD=11).

**Fig E.**
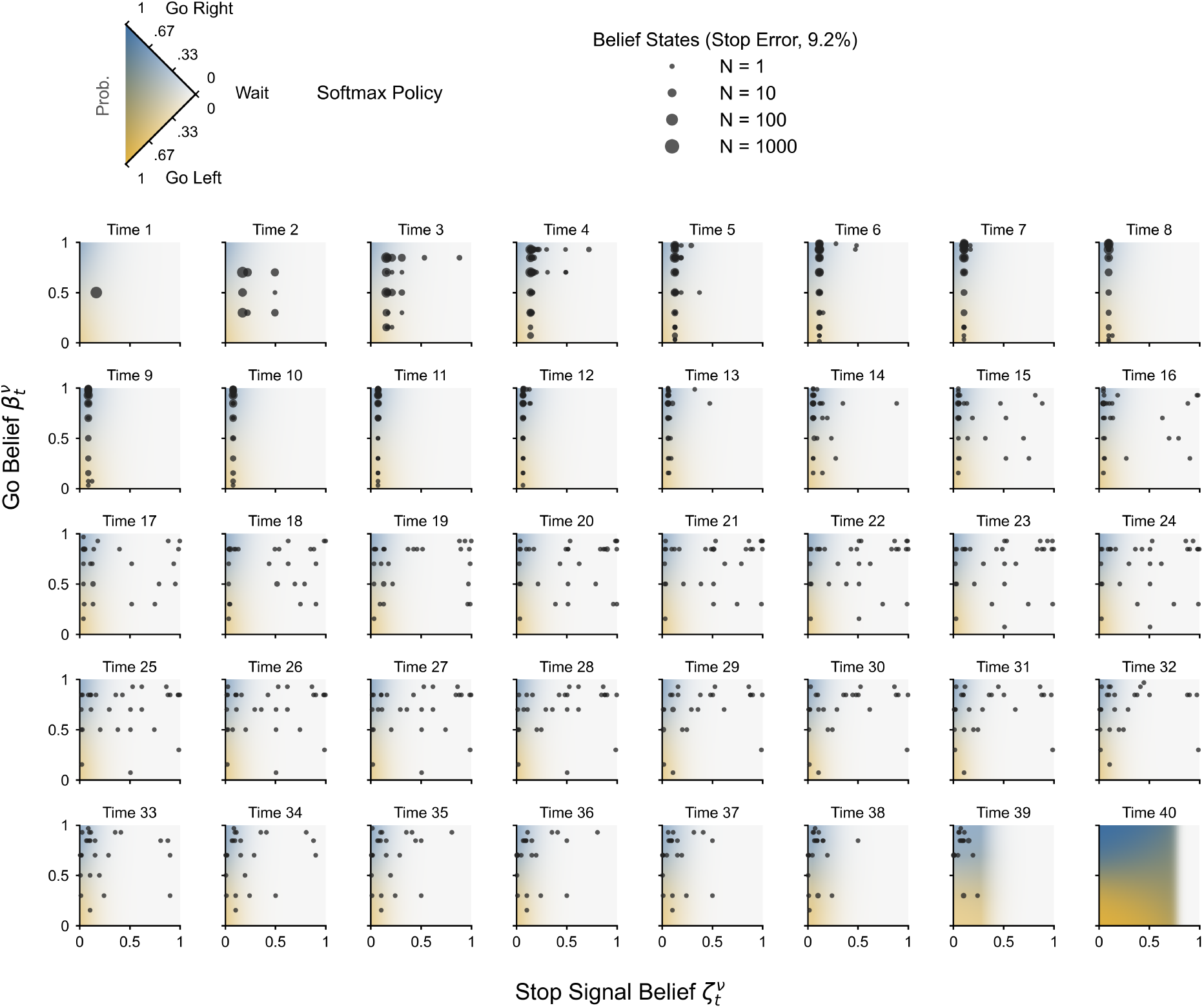
Softmax policy and belief states for Stop Error trials. Background heatmaps represent action probabilities from the softmax policy. Scatter points show simulated trajectories from 1, 200 stop trials (right go cue, SSD=11).

### A.6 Stop prior sensitivity analysis

In the ABCD task, stop trials account for only 1/6 of the total trials (60/360), which limits the natural fluctuation of participants’ across-trial stop prior prediction 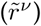. As shown in Fig F, simulating the POMDP model across a realistic range of prior probabilities (0.05 to 0.30) generates small but observable shifts in performance, such as slight increases in Go reaction times, as well as in Stop trial accuracy and error reaction times. Given this restricted range, we fixed 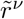 to 1/6 for the current study to simplify the model and focus on estimating within-trial dynamics. This step allows us to establish stable within-trial parameters first, which can then serve as a baseline for future work to free the stop prior and investigate across-trial learning effects.

**Fig F.**
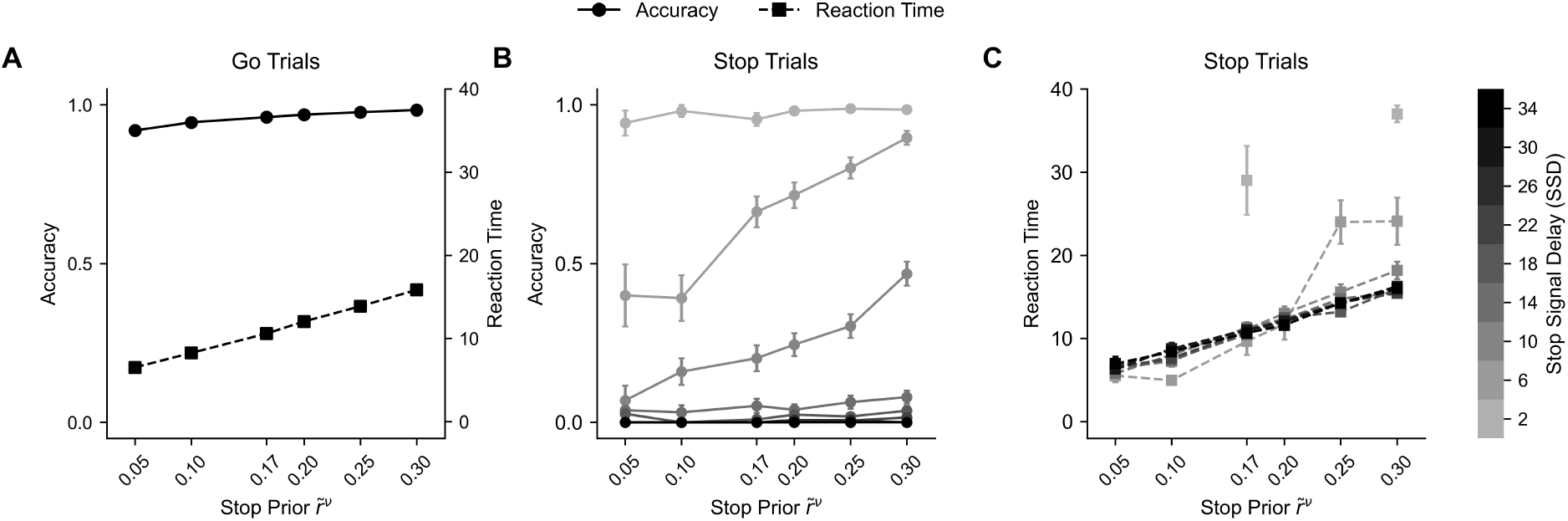
Sensitivity of POMDP behavioral metrics to stop trial priors. Simulated outputs across varying predicted stop prior 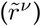, centered around the ABCD SST baseline stop rate (1/6). (A) Go accuracy (solid/circles; left axis) and reaction times (dashed/squares; right axis). (B-C) Stop accuracy and reaction times across Stop Signal Delays (SSD: 2–34, grayscale). Error bars denote standard error of the mean (s.e.m.).

## B TeSBI architecture and validation

### B.1 TeSBI architecture

The TeSBI (Transformer-encoded Simulation-Based Inference) pipeline employs an end-to-end architecture, jointly optimizing a Transformer encoder and a Neural Spline Flow density estimator to perform amortized inference of subject-level parameter posteriors.

The overall network architecture is illustrated in Fig G. The procedure for both the amortized training phase and the posterior inference phase is detailed in Algorithm 1.

**Fig G.**
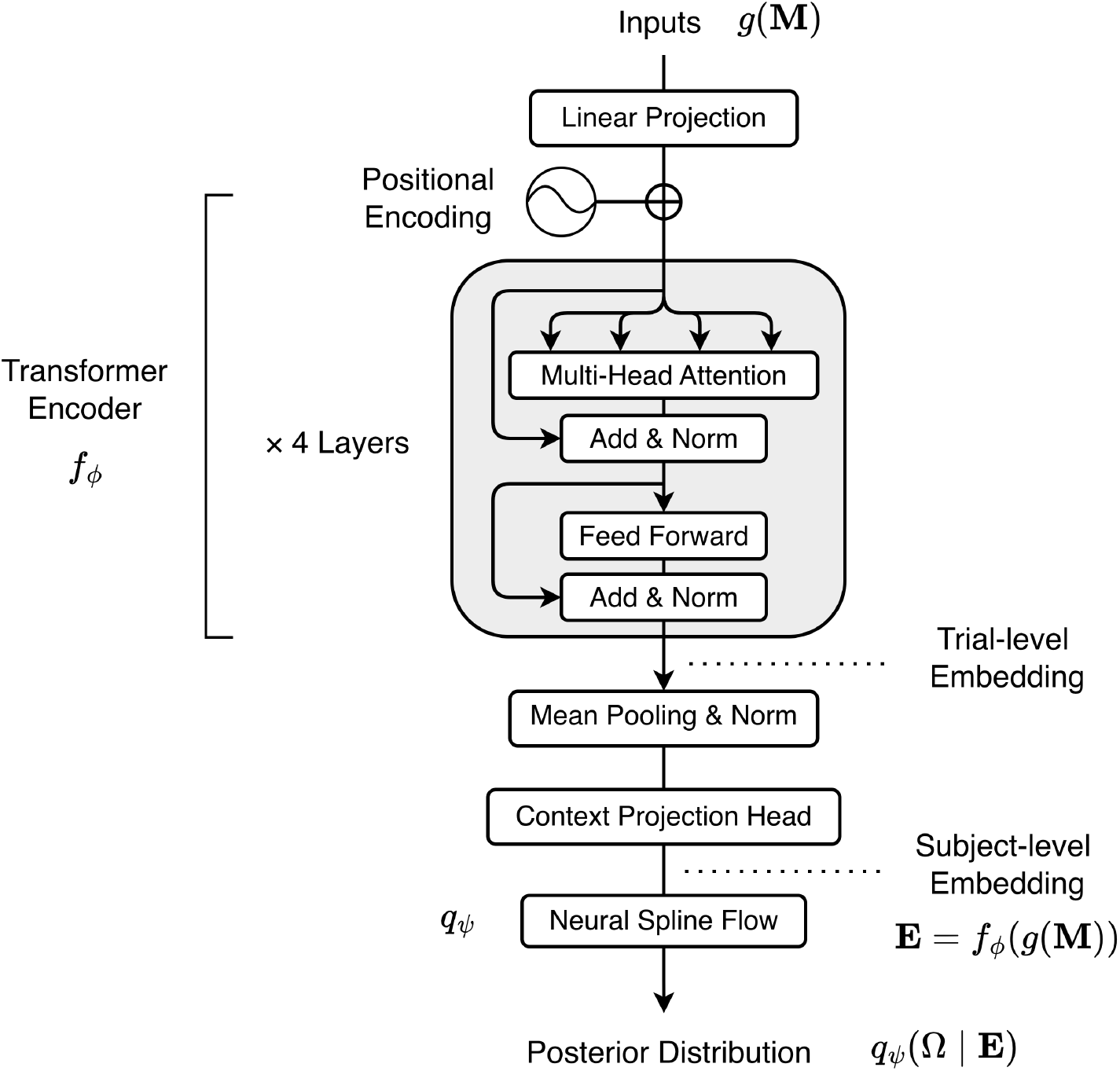
TeSBI architecture. The input feature sequence *g*(**M**) is linearly projected and processed by a 4-layer Transformer encoder *f*_*ϕ*_ to extract trial-level embeddings. These embeddings are aggregated via global mean pooling and refined by a context projection head to yield the subject-level summary embedding **E**. Finally, this embedding directly conditions a Neural Spline Flow density estimator to approximate the subject-level posterior distribution *q*_*Ψ*_(Ω | **E**) of the POMDP parameters.

#### Algorithm 1

Transformer-encoded Simulation-Based Inference (TeSBI)

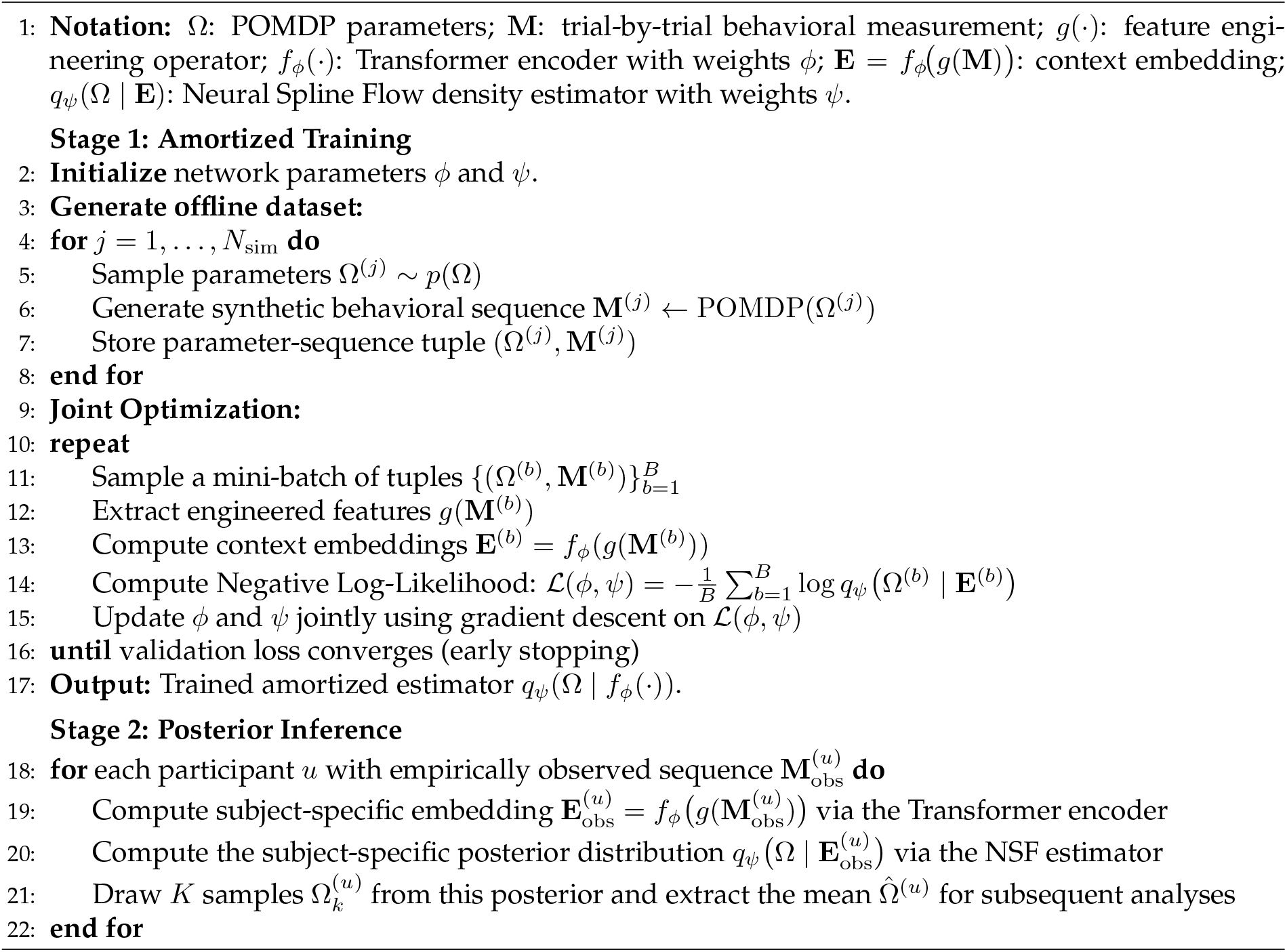

#### Implementation details

The end-to-end TeSBI pipeline was implemented in PyTorch, utilizing the zuko library (v1.6.0) for the normalizing flow estimation.

The 18-dimensional trial-level features were first linearly projected into a continuous latent space. The Transformer encoder then operated on this projected model dimension of 128 with sinusoidal positional encodings. It comprised 4 layers with 8 attention heads each, and a feedforward dimension of 512. Trial-level representations were aggregated via global mean pooling over the temporal dimension, followed by a context projection head (comprising layer normalization, a linear projection, ReLU activation, and a final linear layer) to yield a 128-dimensional summary embedding. For density estimation, we employed a Neural Spline Flow conditioned directly on this embedding, utilizing 2 hidden layers with 256 features each.

Amortized training was performed on a dataset of *N*_sim_ = 100,000 simulated behavioral sessions, partitioned into an 80% training set and a 20% validation set. The unified network jointly minimized the negative log-likelihood of the ground-truth parameters given the behavioral sequences. Optimization was conducted using the AdamW optimizer (batch size 128, initial learning rate 10^*™*3^, weight decay 10^*™*5^). To ensure stable optimization across the joint architecture, gradients were clipped to a maximum norm of 5.0. Furthermore, we employed a learning rate scheduler (reducing the learning rate by a factor of 0.5 upon a validation loss plateau of 5 epochs) and an early stopping mechanism (with a patience of 15 epochs) to prevent overfitting.

### B.2 Transformer encoded representations

Our Transformer encoder compresses the sequential data from 360 trials into a 128-dimensional embedding vector, **E** = *f*_*ϕ*_ *g*(**M**). This approach maps the complex, dynamic behavioral profile of each participant onto a single point in a high-dimensional latent space (R^128^). Fig H shows the Pearson correlation coefficients between model-agnostic behavioral metrics and the latent embeddings, showing some strong relationships.

To examine the validity of the learned representations, we perform a Canonical Correlation Analysis (CCA) between the 128-dimensional latent embeddings and 15 extracted model-agnostic behavioral metrics. After visualizing the initial pairwise correlations in Fig HA, a permutation test (1000 iterations) reveals that all 15 canonical variates are statistically significant (all *p <* 0.001, with canonical correlations *ρ*_*c*_ = 0.995, 0.983, 0.928, 0.868, 0.693, 0.625, 0.527, 0.415, 0.335, 0.287, 0.263, 0.209, 0.192, 0.179, and 0.169).

As shown in Figs HB and HC, the first canonical variate captures a substantial correlation (*ρ*_*c*_ = 0.99) and reveals how both static and dynamical behaviors are holistically encoded across the latent space, confirming the strong expressive capacity of the TeSBI embeddings.

**Fig H.**
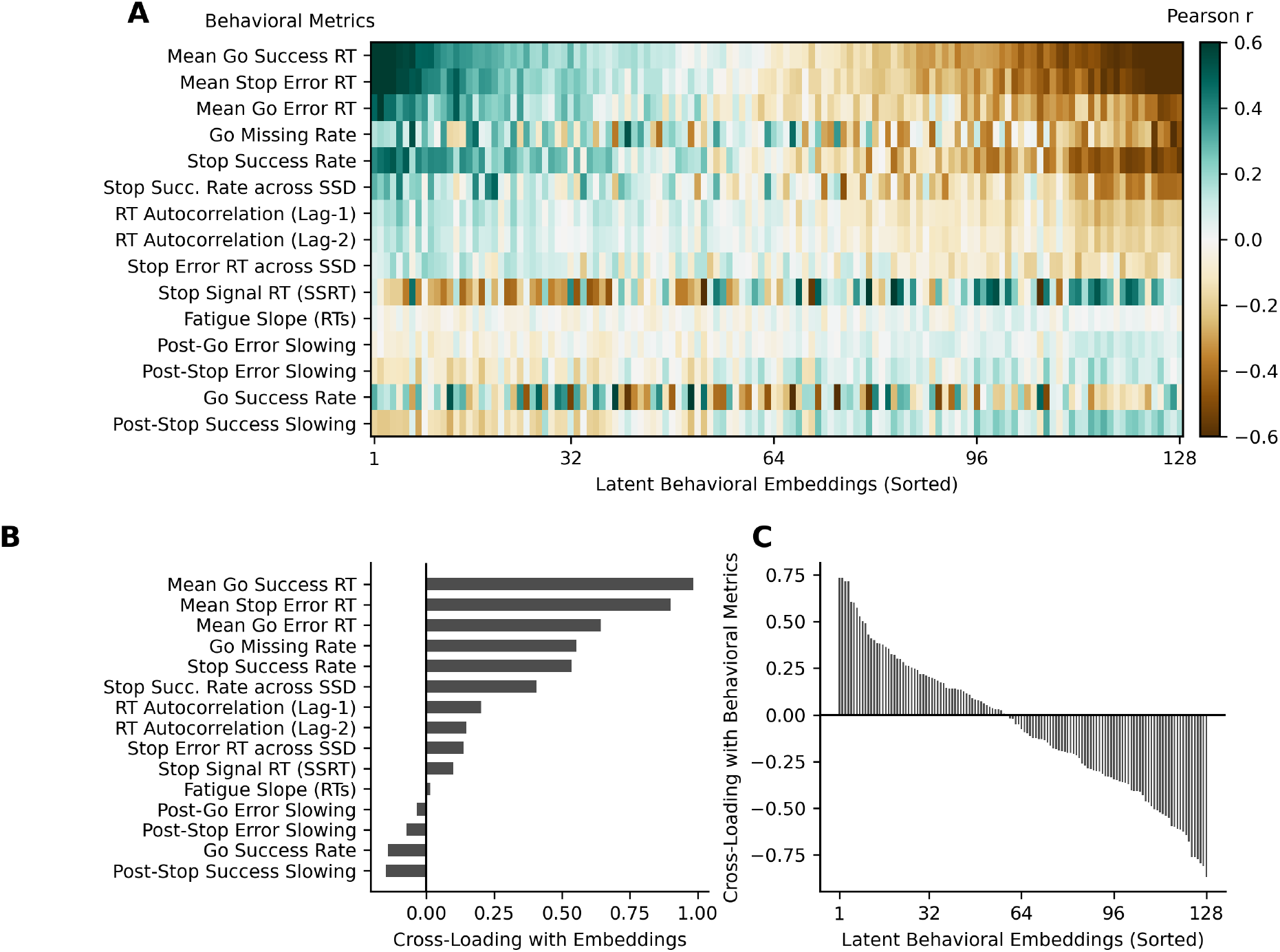
Canonical Correlation Analysis linking learned embeddings and behavioral metrics. (A) Heatmap illustrating the pairwise Pearson correlation coefficients between the 128 latent embeddings (x-axis, sorted by canonical loadings) and the 15 extracted behavioral metrics (y-axis). (B) Sorted cross-loadings of the behavioral metrics on the first canonical variate (*ρ*_*c*_ = 0.995). The encoder effectively captures both static task performance (e.g., mean Go RT, SSRT) and temporal dynamics (e.g., post-stop slowing, RT autocorrelation). (C) Sorted cross-loadings of the latent embeddings on the first canonical variate. The graded distribution indicates that behavioral information is encoded holistically across the latent space rather than being localized to a few dominant dimensions. Only the first canonical variate that capturing the largest shared variance is visualized in panels B and C to avoid visual clutter.

To visualize the structure of these behavioral fingerprints, we apply Principal Component Analysis (PCA) to project the latent embeddings into 2D space (R^2^), capturing 39.8% of the total variance (PC 1: 24.7%, PC 2: 17.0%). As shown in Fig I, the distribution of participant embeddings forms a continuous manifold without discrete boundaries. Critically, when mapping clinical scores onto the 2D space, we observe that participants with higher ADHD scores are dispersed throughout the manifold rather than forming a distinct or isolated “disorder cluster”.

This dispersion provides visual confirmation of our regression results: higher ADHD scores are not linked to a single, unique computational failure factor. Instead, higher ADHD scores are compatible with diverse computational profiles, ranging from varied sensory processing capacities to differing cost valuations. This heterogeneity supports dimensional frameworks (e.g., RDoC) that conceptualize clinical labels as broad, overlapping distributions along a continuous cognitive spectrum, rather than categorical deficits defined by a single phenotype.

**Fig I.**
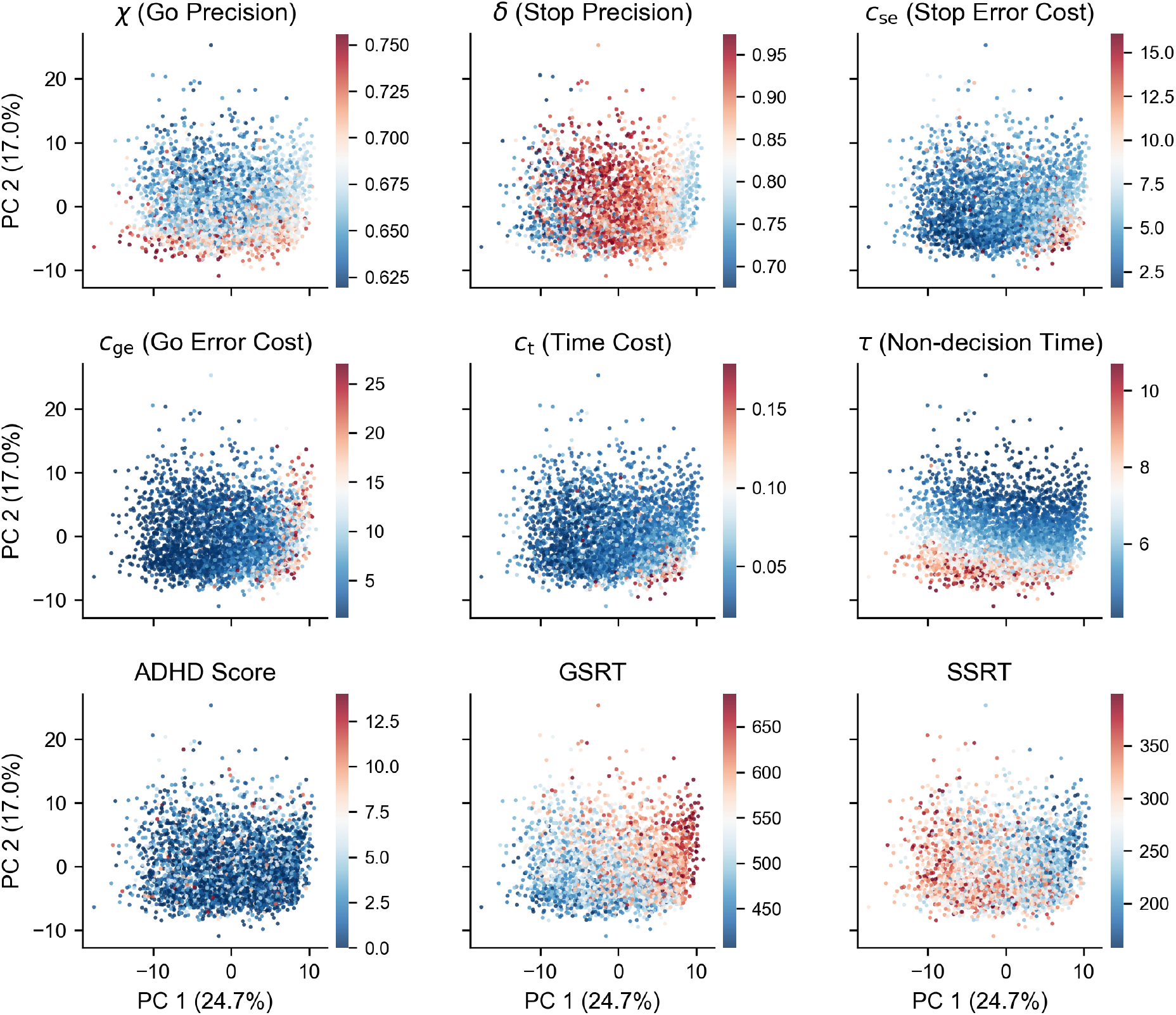
2D PCA projection of the behavioral embedding space. Participant embeddings are projected onto a 2D space using PCA (explaining 41.7% of total variance). Each point represents a single participant (*N* = 3, 567). Panels are colored by POMDP parameters and behavioral metrics to visualize the continuous nature of the latent behavioral landscape. Notably, participants with higher ADHD scores are heterogeneously distributed throughout the space rather than clustering in a specific region. This illustrates that clinical characteristics map onto a diverse spectrum of computational attributes rather than a single isolated deficit.

## C POMDP model selection and validation

To ensure parameter identifiability while preserving predictive power, we apply an iterative step-down procedure across 15 POMDP variants (Table B). Then, starting from the 10-parameter full model (M10.1), we iteratively reduce unidentifiable parameters. At each step, the parameters with the lowest recovery correlations are fixed to specific constants based on posterior quantiles (20%, 50%, or 80%).

### C.1 Model specification

Table B outlines the specifications and parameter constraints across all variants. The full model (M10.1) includes ten parameters: go null (*χ*^*′*^), go precision (*χ*), stop null (***δ***^*′*^), stop precision (***δ***), go error cost (*c*_ge_), go missing cost (*c*_gm_), stop error cost (*c*_se_), time cost (*c*_*t*_), inverse temperature (*φ*), and non-decision time (*τ*).

The *Prior Range* row defines the estimation boundaries for each parameter. In the subsequent model rows, checkmarks (✓) denote free parameters estimated within these prior ranges, whereas single numerical values indicate that the parameter is fixed to a specific constant.

**Table B.** POMDP model specification and parameter priors.

| Model | $\chi'$ | $\chi$ | $\delta'$ | $\delta$ | $c_{ge}$ | $c_{gm}$ | $c_{se}$ | $c_t$ | $\varphi$ | $\tau$ | $\hat{r}^\nu$ |
| --- | --- | --- | --- | --- | --- | --- | --- | --- | --- | --- | --- |
| Prior Range | [0, 1] | [0.5, 1] | [0, 1] | [0.5, 1] | [1, 50] | [1, 50] | [1, 50] | [0.01, 0.5] | [10, 100] | [4, 16] | Fixed |
| **M10.1** | ✓ | ✓ | ✓ | ✓ | ✓ | ✓ | ✓ | ✓ | ✓ | ✓ | **1/6** |
| **M8.1** | ✓ | ✓ | 0.1348 | ✓ | ✓ | ✓ | ✓ | ✓ | 44.6534 | ✓ | **1/6** |
| **M8.2** | ✓ | ✓ | **0.0787** | ✓ | ✓ | ✓ | ✓ | ✓ | **44.6534** | ✓ | **1/6** |
| **M8.3** | ✓ | ✓ | 0.2441 | ✓ | ✓ | ✓ | ✓ | ✓ | 44.6534 | ✓ | **1/6** |
| **M8.4** | ✓ | ✓ | 0.1348 | ✓ | ✓ | ✓ | ✓ | ✓ | 37.9558 | ✓ | **1/6** |
| **M8.5** | ✓ | ✓ | 0.1348 | ✓ | ✓ | ✓ | ✓ | ✓ | 51.8192 | ✓ | **1/6** |
| **M7.1** | ✓ | ✓ | **0.0787** | ✓ | ✓ | 9.5955 | ✓ | ✓ | **44.6534** | ✓ | **1/6** |
| **M7.2** | ✓ | ✓ | **0.0787** | ✓ | ✓ | 13.1139 | ✓ | ✓ | **44.6534** | ✓ | **1/6** |
| **M7.3** | ✓ | ✓ | **0.0787** | ✓ | ✓ | **15.7821** | ✓ | ✓ | **44.6534** | ✓ | **1/6** |
| **M7.4** | 0.1026 | ✓ | **0.0787** | ✓ | ✓ | ✓ | ✓ | ✓ | **44.6534** | ✓ | **1/6** |
| **M7.5** | 0.2361 | ✓ | **0.0787** | ✓ | ✓ | ✓ | ✓ | ✓ | **44.6534** | ✓ | **1/6** |
| **M7.6** | 0.5328 | ✓ | **0.0787** | ✓ | ✓ | ✓ | ✓ | ✓ | **44.6534** | ✓ | **1/6** |
| **M6.1** | **0.0798** | ✓ | **0.0787** | ✓ | ✓ | **15.7821** | ✓ | ✓ | **44.6534** | ✓ | **1/6** |
| **M6.2** | 0.1671 | ✓ | **0.0787** | ✓ | ✓ | **15.7821** | ✓ | ✓ | **44.6534** | ✓ | **1/6** |
| **M6.3** | 0.4295 | ✓ | **0.0787** | ✓ | ✓ | **15.7821** | ✓ | ✓ | **44.6534** | ✓ | **1/6** |

### C.2 Model selection

In order to find a model of the behavior which fits the data well, but does not overfit, we employ a step-down procedure to remove degrees of freedom that are not recoverable. In the first step of this procedure, the inverse temperature (*φ*) and stop null (***δ***^*′*^) are the primary bottlenecks in M10.1, showing the worst recovery (*r* = 0.30 and *r* = 0.34, respectively; Table C). Fixing these two parameters based on the posterior quantiles of M10.1 yields five 8-parameter variants (M8.1–M8.5). We select M8.2 (*φ* = 44.6534, ***δ***^*′*^ = 0.0787) for the next step due to its superior in-sample (Table D) and out-of-sample (Table E) posterior predictive checks.

In the second step, we address the least recoverable parameters in M8.2 by creating six 7-parameter variants. We fix either the go missing cost (*c*_gm_) (M7.1–M7.3) or the go null parameter (*χ*^*′*^) (M7.4–M7.6) to the posterior quantiles of M8.2. From these, M7.3 (*c*_gm_ = 15.7821) emerges as the best variant.

In the final step, because *χ*^*′*^ remains the sole bottleneck in M7.3 (r = 0.6681), we further reduce the architecture to 6-parameter variants (M6.1–M6.3) by fixing *χ*^*′*^ to the posterior quantiles of M7.3. All M6.1, M6.2 and M6.3 successfully pass the parameter recovery test for all remaining free parameters (Pearson’s *r >* 0.65). Ultimately, we select M6.1 (*c*_gm_ = 2.2687) as the primary computational model. It achieves complete parameter identifiability while maintaining strong in-sample and out-of-sample predictive power (total distances of 1.88611 and 3.77383).

### C.3 Parameter recovery analysis

To evaluate parameter identifiability, we simulate behavior using parameters sampled from the priors, fit the models to these simulated datasets, and calculate the Pearson correlation coefficient (*r*) between the ground-truth parameters and the recovered estimates.

To ensure robustness, we apply a criterion requiring *r >* 0.65 for all estimated parameters (indicated by checkmarks in the Valid column). As Table C shows, while increasing the number of free parameters enhances model flexibility to fit the data, it also introduces severe parameter unidentifiability. Our step-down procedure effectively resolves this issue. In the final selected model (M6.1), all six free parameters strictly satisfy this identifiability criterion: go precision (*r* = 0.89), stop precision (*r* = 0.72), non-decision time (*r* = 0.83), stop error cost (*r* = 0.81), time cost (*r* = 0.82), and go error cost (*r* = 0.72).

**Table C.** Parameter recovery correlations across model variants.

| Model | $\chi$ | $\delta$ | $\tau$ | $c_{se}$ | $c_t$ | $c_{ge}$ | $c_{gm}$ | $\chi'$ | $\delta'$ | $\varphi$ | Valid |
| --- | --- | --- | --- | --- | --- | --- | --- | --- | --- | --- | --- |
| M10.1 | 0.7338 | 0.52576 | 0.76639 | 0.71686 | 0.77255 | 0.67475 | 0.25241 | 0.68363 | 0.28484 | 0.23538 |  |
| M8.1 | 0.78803 | 0.67084 | 0.76253 | 0.72547 | 0.64495 | 0.65898 | 0.19468 | 0.56529 | — | — |  |
| M8.2 | 0.75792 | 0.63352 | 0.75234 | 0.72376 | 0.68564 | 0.6542 | 0.18572 | 0.58766 | — | — |  |
| M8.3 | 0.78363 | 0.65966 | 0.75261 | 0.74255 | 0.69466 | 0.65715 | 0.25048 | 0.5983 | — | — |  |
| M8.4 | 0.77669 | 0.67832 | 0.78664 | 0.7536 | 0.66689 | 0.65421 | 0.20505 | 0.58863 | — | — |  |
| M8.5 | 0.76724 | 0.62577 | 0.7137 | 0.73714 | 0.67105 | 0.63331 | 0.26696 | 0.58201 | — | — |  |
| M7.1 | 0.77864 | 0.73923 | 0.78226 | 0.8094 | 0.74747 | 0.56983 | — | 0.66981 | — | — |  |
| M7.2 | 0.74035 | 0.74885 | 0.80225 | 0.82987 | 0.76128 | 0.60295 | — | 0.66004 | — | — |  |
| M7.3 | 0.77628 | 0.75358 | 0.78879 | 0.82022 | 0.7603 | 0.6681 | — | 0.57688 | — | — |  |
| M7.4 | 0.78577 | 0.68973 | 0.69869 | 0.81331 | 0.73715 | 0.59321 | 0.24727 | — | — | — |  |
| M7.5 | 0.80723 | 0.70444 | 0.71446 | 0.85211 | 0.78155 | 0.61792 | 0.29937 | — | — | — |  |
| M7.6 | 0.78866 | 0.70419 | 0.70713 | 0.79195 | 0.76916 | 0.57304 | 0.19798 | — | — | — |  |
| **M6.1** | **0.89382** | **0.72778** | **0.83096** | **0.81072** | **0.82478** | **0.72995** | — | — | — | — | ✓ |
| M6.2 | 0.88438 | 0.69043 | 0.84585 | 0.80845 | 0.80418 | 0.71801 | — | — | — | — | ✓ |
| M6.3 | 0.89813 | 0.70072 | 0.81195 | 0.76931 | 0.76785 | 0.70143 | — | — | — | — | ✓ |

### C.4 Posterior predictive checks

To validate the predictive power of the models, we perform in-sample posterior predictive checks (PPCs) by simulating the entire task behavior for each participant using their posterior mean estimates. We then quantify the empirical distance between the simulated and observed data across multiple metrics: choice proportions (percentages of Go Success, Go Error, Go Missing, and Stop Success) and reaction time distributions (measured via Wasserstein and Kolmogorov-Smirnov distances for Go Success and Stop Error trials).

To evaluate generalization and prevent overfitting, we further apply an out-of-sample temporal crossvalidation PPC. For each participant, we use the first 80% of trials to estimate the posterior parameters and simulate the subsequent behavior. We then compute the empirical distances strictly on the withheld 20% of trials, across the same metrics.

The empirical distances for the in-sample and out-of-sample checks are summarized in Table D and Table E, respectively.

**Table D.** In-sample empirical distances from posterior predictive checks.

| Model | Choice Proportions |  |  |  | WS Dist. (RT) |  |  | KS Dist. (RT) |  | Total Dist. |
| --- | --- | --- | --- | --- | --- | --- | --- | --- | --- | --- |
|  | GS | GE | GM | SS | GS | GE | SE | GS | SE |  |
| M10.1 | 0.02235 | 0.0096 | 0.01598 | 0.01478 | 0.02312 | 0.17258 | 0.07416 | 0.09356 | 0.2337 | 2.26267 |
| M8.1 | 0.019 | 0.00756 | 0.0153 | 0.01493 | 0.02435 | 0.14736 | 0.05723 | 0.10057 | 0.18695 | 2.28152 |
| M8.2 | 0.0214 | 0.00885 | 0.01558 | 0.01413 | 0.0191 | 0.15994 | 0.04766 | 0.0786 | 0.16869 | 1.97597 |
| M8.3 | 0.01516 | 0.0069 | 0.01387 | 0.02005 | 0.02181 | 0.15718 | 0.09139 | 0.09118 | 0.24394 | 2.48218 |
| M8.4 | 0.01738 | 0.00567 | 0.01585 | 0.01493 | 0.02061 | 0.15326 | 0.05907 | 0.08493 | 0.18203 | 2.17317 |
| M8.5 | 0.01908 | 0.00778 | 0.01507 | 0.01431 | 0.01819 | 0.16187 | 0.06066 | 0.07338 | 0.18544 | 2.0013 |
| M7.1 | 0.02158 | 0.00928 | 0.01622 | 0.01534 | 0.0264 | 0.14367 | 0.05022 | 0.10922 | 0.18653 | 2.26475 |
| M7.2 | 0.01928 | 0.00829 | 0.01518 | 0.01438 | 0.02214 | 0.1519 | 0.04803 | 0.08974 | 0.17741 | 1.87367 |
| M7.3 | 0.01744 | 0.00767 | 0.01413 | 0.01392 | 0.0219 | 0.15599 | 0.05317 | 0.08815 | 0.17837 | 1.86933 |
| M7.4 | 0.01857 | 0.00806 | 0.01624 | 0.01407 | 0.02187 | 0.14429 | 0.04924 | 0.08998 | 0.17694 | 1.92229 |
| M7.5 | 0.01791 | 0.00987 | 0.01524 | 0.01486 | 0.01962 | 0.15251 | 0.05456 | 0.07839 | 0.1817 | 2.15058 |
| M7.6 | 0.01528 | 0.00675 | 0.01382 | 0.01672 | 0.02052 | 0.17389 | 0.05394 | 0.08044 | 0.18376 | 2.90014 |
| **M6.1** | **0.02312** | **0.00963** | **0.01679** | **0.01407** | **0.02051** | **0.13895** | **0.04819** | **0.08458** | **0.17723** | **1.88611** |
| M6.2 | 0.01561 | 0.008 | 0.01432 | 0.01518 | 0.01854 | 0.1525 | 0.04877 | 0.07507 | 0.1707 | 2.09818 |
| M6.3 | 0.01323 | 0.00667 | 0.01356 | 0.01617 | 0.01895 | 0.16565 | 0.05624 | 0.07671 | 0.18724 | 2.6225 |

**Table E.** Out-of-sample empirical distances from posterior predictive checks.

| Model | Choice Proportions |  |  |  | WS Dist. (RT) |  |  | KS Dist. (RT) |  | Total Dist. |
| --- | --- | --- | --- | --- | --- | --- | --- | --- | --- | --- |
|  | GS | GE | GM | SS | GS | GE | SE | GS | SE |  |
| M10.1 | 0.0534 | 0.02668 | 0.03917 | 0.07457 | 0.04385 | 0.30011 | 0.08903 | 0.16962 | 0.35948 | 4.13818 |
| M8.1 | 0.05204 | 0.02643 | 0.03877 | 0.07455 | 0.04401 | 0.28826 | 0.08392 | 0.168 | 0.35133 | 4.14454 |
| M8.2 | 0.0532 | 0.02668 | 0.03906 | 0.07449 | 0.04225 | 0.29657 | 0.0777 | 0.16371 | 0.34683 | 3.94099 |
| M8.3 | 0.04942 | 0.02606 | 0.03809 | 0.0768 | 0.04415 | 0.29137 | 0.10524 | 0.17038 | 0.3599 | 4.15066 |
| M8.4 | 0.05164 | 0.02636 | 0.03961 | 0.07468 | 0.04289 | 0.29216 | 0.08498 | 0.16388 | 0.34708 | 4.10278 |
| M8.5 | 0.05151 | 0.02621 | 0.03846 | 0.07447 | 0.04297 | 0.29585 | 0.08613 | 0.16441 | 0.3492 | 3.99578 |
| M7.1 | 0.05424 | 0.02745 | 0.04 | 0.07486 | 0.04499 | 0.28682 | 0.08011 | 0.17189 | 0.35604 | 4.15498 |
| M7.2 | 0.05191 | 0.02649 | 0.0383 | 0.07517 | 0.0422 | 0.29192 | 0.07768 | 0.16093 | 0.34967 | 3.93033 |
| M7.3 | 0.05207 | 0.02603 | 0.03922 | 0.07472 | 0.04323 | 0.29094 | 0.08061 | 0.16571 | 0.35008 | 3.94844 |
| M7.4 | 0.05151 | 0.02725 | 0.03885 | 0.0754 | 0.04388 | 0.29 | 0.07913 | 0.16718 | 0.35132 | 3.81917 |
| M7.5 | 0.04963 | 0.02709 | 0.03817 | 0.07459 | 0.04262 | 0.29247 | 0.08094 | 0.16392 | 0.34849 | 3.95992 |
| M7.6 | 0.05061 | 0.02589 | 0.03811 | 0.07401 | 0.04299 | 0.30578 | 0.08067 | 0.16337 | 0.34761 | 4.8208 |
| **M6.1** | **0.05542** | **0.02753** | **0.04007** | **0.07639** | **0.04311** | **0.28142** | **0.08023** | **0.16364** | **0.34904** | **3.77383** |
| M6.2 | 0.05014 | 0.02768 | 0.03854 | 0.07682 | 0.0432 | 0.29928 | 0.08049 | 0.16249 | 0.3459 | 3.88856 |
| M6.3 | 0.04648 | 0.02659 | 0.03734 | 0.0746 | 0.04311 | 0.30706 | 0.08406 | 0.16387 | 0.34985 | 4.57465 |

## D Comparison with alternative models

While the POMDP and RDEX models (Weigard et al., 2023) offer distinct mechanistic explanations for response inhibition, dynamic belief updating and subjective cost evaluation (POMDP) versus an accumulate-to-threshold process (RDEX), both generate comparable simulated behavior. To evaluate this, we performed a posterior predictive check comparing our POMDP framework against the best-performing RDEX variant.

To generate the simulated data, we selected three representative participants corresponding to the 95th, 50th, and 5th percentiles of the POMDP model goodness-of-fit distribution, simulating 5 task sessions per participant. For the RDEX model, we utilized the parameters for these same subjects estimated via DE-MCMC, as described in Weigard et al. (2023).

As shown in Fig J, both models successfully capture the core behavioral signatures of the task, providing adequate and comparable fits to the empirical observations. Specifically, both simulations reproduce the global outcome proportions, the right-skewed reaction time distributions for Go Success and Stop Error trials, the trial-by-trial tracking of the SSD staircases, and the inhibition performance across SSDs.

**Fig J.**
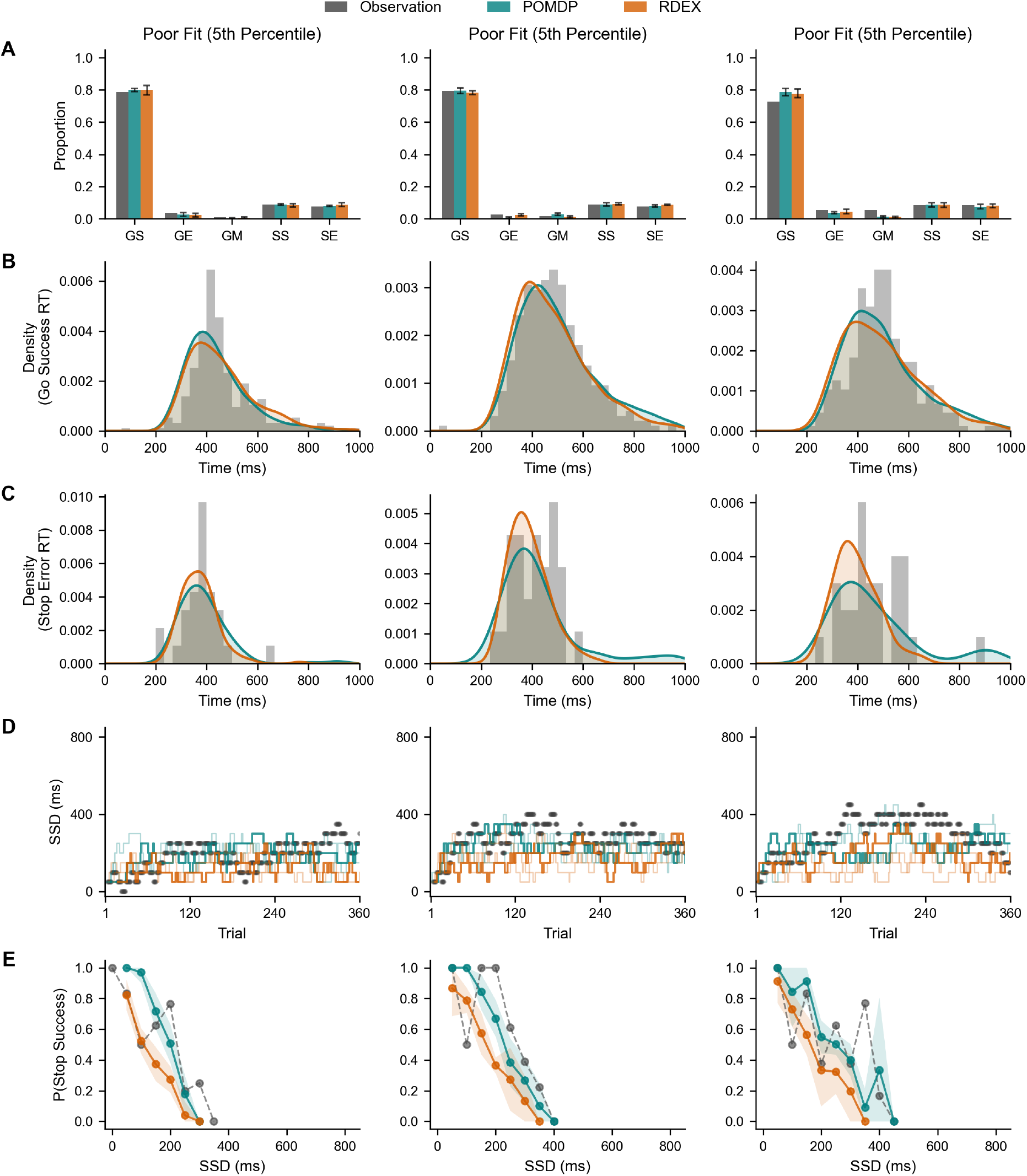
Qualitative comparison of the POMDP and RDEX models. The columns represent three representative participants corresponding to the 95th, 50th, and 5th percentiles of the POMDP model fit distribution. The panels display simulated versus observed data across five key behavioral metrics: (A) outcome proportions, reaction time distributions for (B) Go Success and (C) Stop Error trials, (D) SSD staircase trajectories, and (E) stop accuracy across SSD. Error bars and shaded regions represent the s.e.m. Overall, although mechanistically different, both the POMDP and RDEX models yield comparable and adequate fits to the empirical data.

## E Computational attributes

### E.1 Aggregate distributions of computational attributes

Across the entire baseline cohort, the estimated parameters exhibit distinct aggregate profiles. The sensory parameters, go precision (*χ*) and stop precision (***δ***), are both tightly clustered. This indicates high perceptual consistency across the population, although ***δ*** features a slightly longer left tail, reflecting greater variability in stop signal processing.

In contrast, all three intrinsic cost parameters (*c*_se_, *c*_ge_, and *c*_*t*_) exhibit distinct positive skew. This indicates that while most participants operate with relatively low subjective costs for errors and time delays, a long tail of individuals heavily penalizes these suboptimal outcomes. Finally, the non-decision time (*τ*) also exhibits a right-skewed distribution, averaging 5.7 steps (approximately 142 ms), which aligns well with known physiological bounds for visuo-motor delays.

**Fig K.**
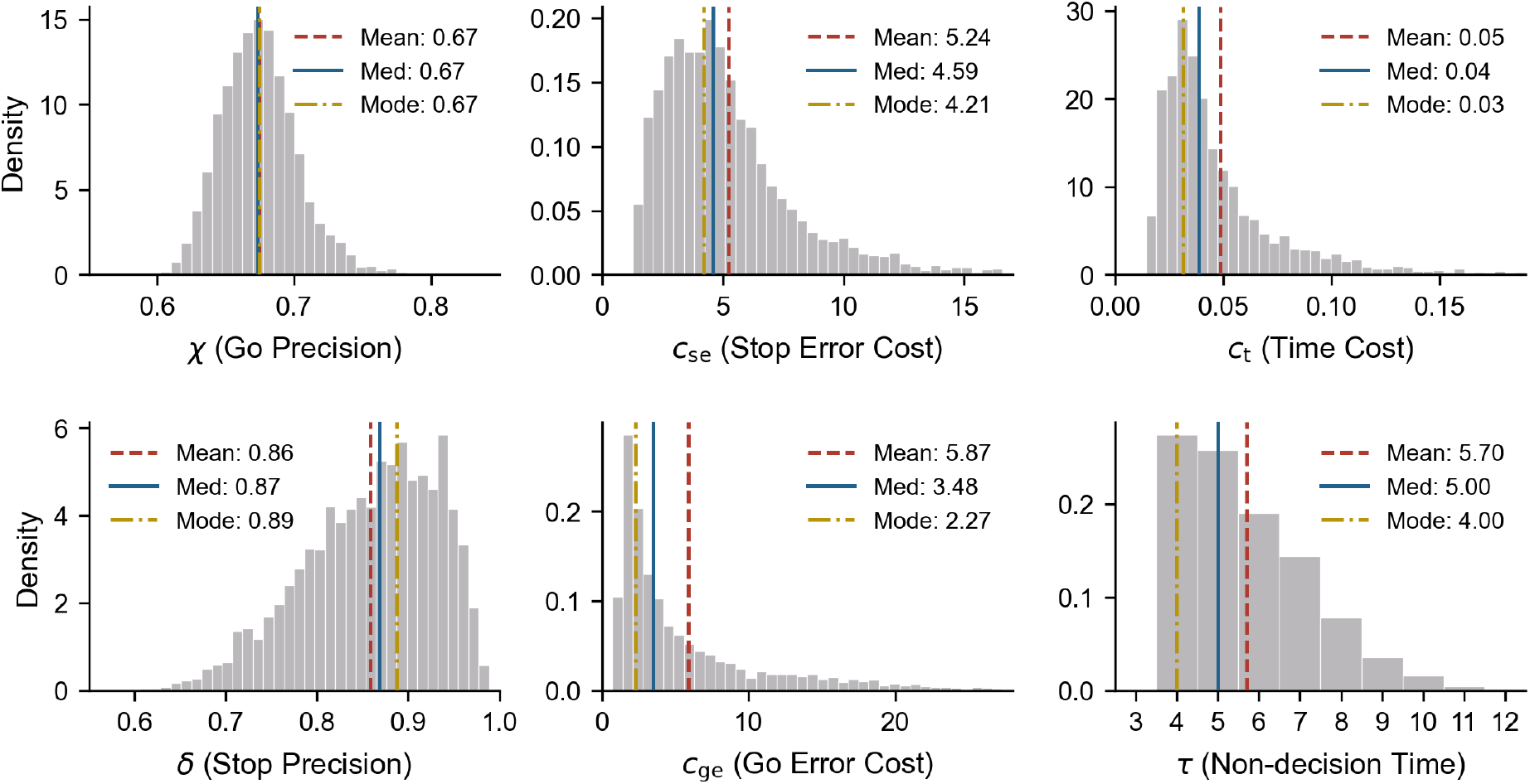
Aggregate distributions of participant-level computational attributes. Distributions of individual point estimates across all participants (*N* = 3, 567). For each participant, the point estimate is defined as the mean of 1,000 samples drawn from their respective posterior distribution. The parameters exhibit distinct distributional shapes: the sensory parameters (*χ*, ***δ***) are tightly clustered, while the non-decision time (*τ*) and all three cost parameters (*c*_se_, *c*_ge_, *c*_*t*_) display distinct positive skew. Summary statistics are provided as Mean / Median / Mode: *χ* = 0.67*/*0.67*/*0.67; ***δ*** = 0.86*/*0.87*/*0.89; *c*_se_ = 5.24*/*4.59*/*4.21; *c*_ge_ = 5.87*/*3.48*/*2.27; *c*_*t*_ = 0.05*/*0.04*/*0.03; *τ* = 5.70*/*5.00*/*4.00.

### E.2 Parameter regression and interaction analysis

To evaluate the association between ADHD scores and the estimated POMDP parameters systematically while accounting for potential confounding factors, we perform multiple linear regression analyses. For each computational parameter Ω ∈ {*χ*, ***δ***, *c*_se_, *c*_ge_, *c*_t_, *τ}*, the baseline main-effects model is formulated as follows:

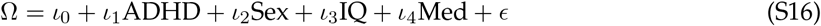

To ensure that the associations described in the main text are robust and consistent across demographic or clinical profiles, we extend this baseline model to include interaction effects. Specifically, for each parameter, we independently add two-way interaction terms (ADHD *×* Sex, ADHD *×* IQ, and ADHD × Medication) to the main model. We also evaluate a full model that simultaneously incorporates all three interaction terms.

Statistical model parsimony and goodness-of-fit are assessed using the Bayesian Information Criterion (BIC). As detailed in Table F, the main-effects baseline model consistently generates the lowest BIC (ΔBIC = 0) across all six parameters. The inclusion of most individual interaction terms increases the BIC substantially (ΔBIC *>* 4), with the minor exception of the ADHD *×* Medication interaction on stop precision (***δ***; ΔBIC = 1.7). Furthermore, the full interaction models are heavily penalized (ΔBIC ≈16–24), indicating strong evidence against the added complexity.

The inclusion of interaction terms for demographic and clinical variables does not produce a significant improvement in model fit. Consequently, these interaction terms are excluded, and the main-effects model is retained as the most parsimonious and optimal. Detailed results elucidating the specific relationships between these computational attributes and clinical scores under the selected main-effects model are presented in the main text.

**Table F.**
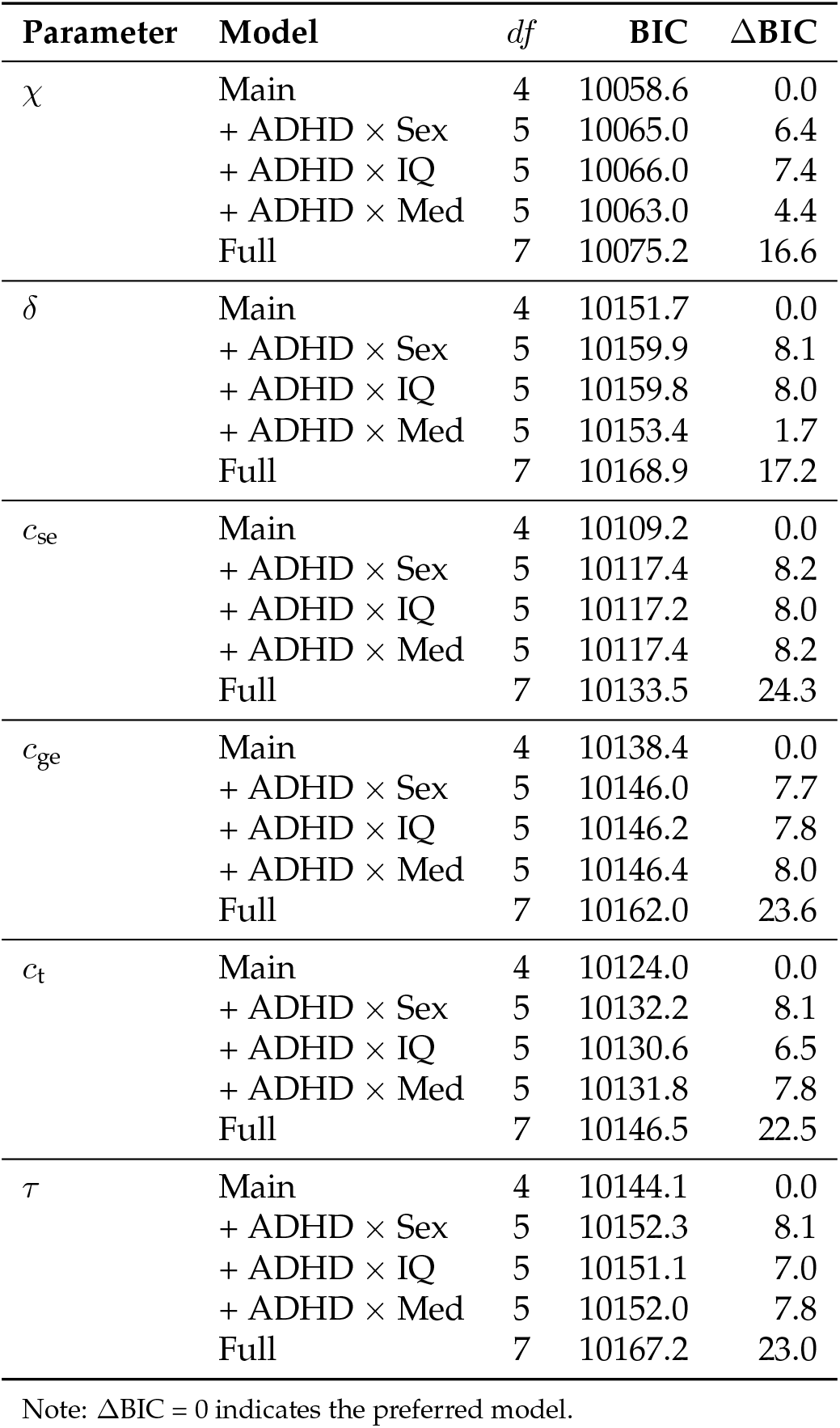
Supplementary: BIC Model Comparisons.

### E.3 Predictor sets comparison

To evaluate the predictive performance of the computational attributes relative to classic behavioral metrics in predicting ADHD scores, we compare three distinct predictor sets: (1) Classic metrics: SSRT, mean Go reaction time (Mean Go RT), and standard deviation of Go reaction time (Go RT SD). Notably, SSRT is derived by subtracting the mean SSD from the mean Go RT, which might be biased due to violations of the independence assumption; (2) POMDP Model: the six computational attributes; and 3) Combination: a combination of both sets. All regression models include sex, IQ, and medication status as baseline covariates.

As summarized in Table G, the POMDP Model explains slightly more variance (Adjusted *R*^2^ = 0.1768) than the Classic metrics (Adjusted *R*^2^ = 0.1759). Furthermore, a Likelihood Ratio Test (LRT) comparing the Combination model against the baseline confirms that incorporating the POMDP parameters significantly improves the overall model fit (*p* = 0.026). This confirms that, beyond within-trial cognitive dynamics, the computational attributes capture unique, clinically relevant variance. Despite the BIC penalty for added complexity (BIC: 9495.2 vs. 9477.8), the favorable AIC (AIC: 9433.4 vs. 9434.5) and significant LRT highlight the POMDP model’s superior explanatory power for ADHD.

**Table G.** Regression predictor set comparison.

| Predictor Set | Variables | Adjusted $R^2$ | AIC | BIC | LRT $p$-value |
| --- | --- | --- | --- | --- | --- |
| Classic metrics | SSRT, Go RT Mean, Go RT SD | 0.1759 | 9434.5 | 9477.8 | — |
| POMDP Model | POMDP Parameters | 0.1768 | 9433.4 | 9495.2 | — |
| Combination | Classic metrics + POMDP Parameters | 0.1778 | 9432.1 | 9512.4 | 0.026 |

## F Model-agnostic statistical analysis

Table H shows the number of subjects excluded by each quality control criterion. Importantly, the exclusion counts listed for each criterion are independent; a single subject may fail multiple criteria simultaneously. After applying all filters of experimental quality, behavioral performance, and demographic data completeness, the final valid dataset includes 3,567 baseline subjects (31.0% of the original sample).

**Table H.**
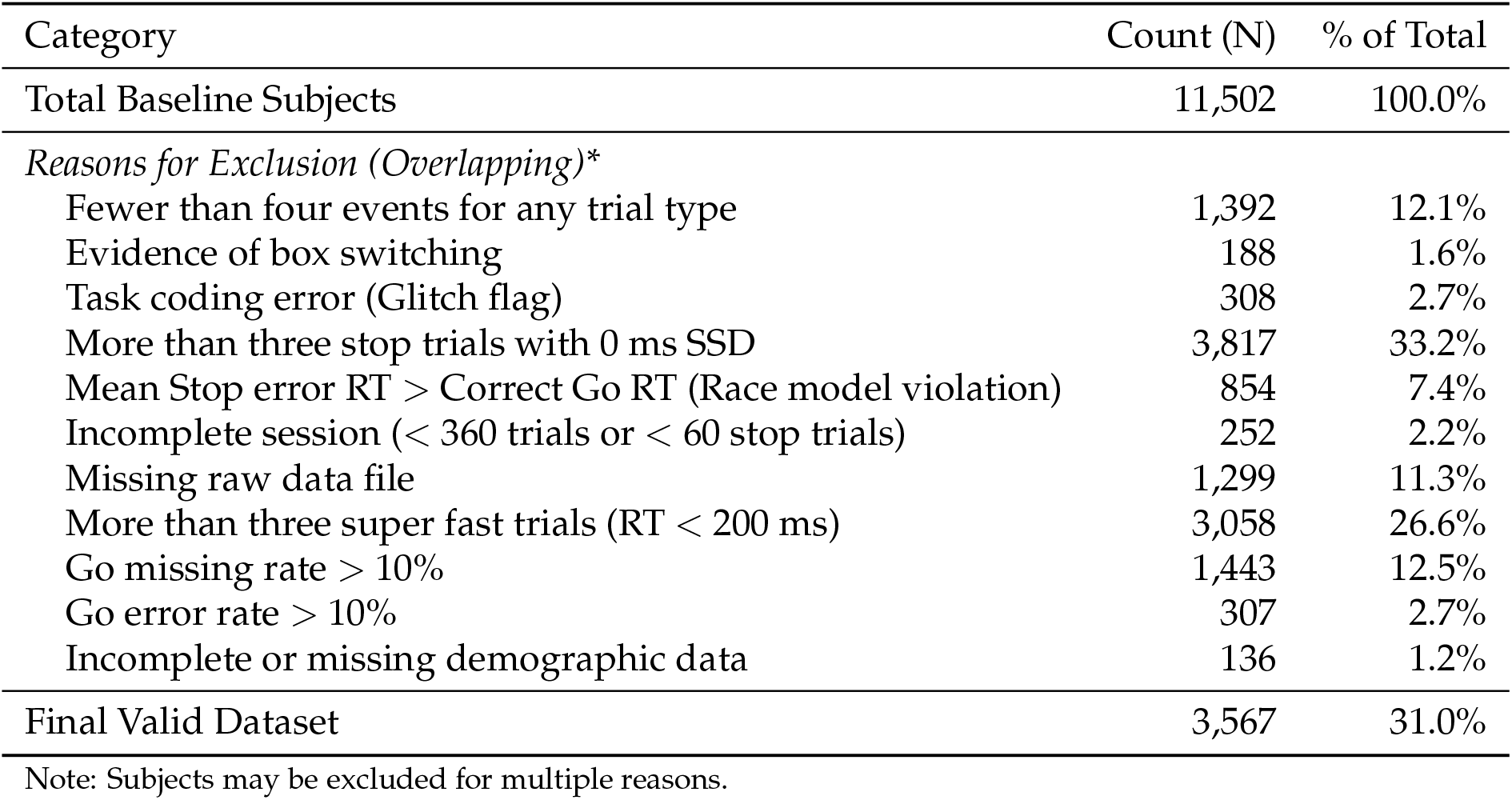
Subject inclusion and exclusion criteria.

| Category | Count (N) | % of Total |
| --- | --- | --- |
| Total Baseline Subjects | 11,502 | 100.0% |
| *Reasons for Exclusion (Overlapping)\** |  |  |
| Fewer than four events for any trial type | 1,392 | 12.1% |
| Evidence of box switching | 188 | 1.6% |
| Task coding error (Glitch flag) | 308 | 2.7% |
| More than three stop trials with 0 ms SSD | 3,817 | 33.2% |
| Mean Stop error RT > Correct Go RT (Race model violation) | 854 | 7.4% |
| Incomplete session (< 360 trials or < 60 stop trials) | 252 | 2.2% |
| Missing raw data file | 1,299 | 11.3% |
| More than three super fast trials (RT < 200 ms) | 3,058 | 26.6% |
| Go missing rate > 10% | 1,443 | 12.5% |
| Go error rate > 10% | 307 | 2.7% |
| Incomplete or missing demographic data | 136 | 1.2% |
| Final Valid Dataset | 3,567 | 31.0% |

Note: Subjects may be excluded for multiple reasons.

Table I outlines the distribution of ADHD scores (symptom severity, raw scores 0–14) across key demographic variables. Because the sample demonstrates inherent imbalances, particularly at higher severity levels, we employ multiple linear regression models controlling for Sex, IQ, and Medication status to rigorously isolate the specific behavioral contributions of ADHD symptoms.

**Table I.** Subgroup counts by Sex, IQ, and Medication across ADHD scores.

| Group | ADHD Score | 0 | 1 | 2 | 3 | 4 | 5 | 6 | 7 | 8 | 9 | 10 | 11 | 12 | 13 | 14 | Total |
| --- | --- | --- | --- | --- | --- | --- | --- | --- | --- | --- | --- | --- | --- | --- | --- | --- | --- |
|  | Count | 1390 | 656 | 373 | 316 | 225 | 206 | 129 | 103 | 57 | 40 | 22 | 20 | 13 | 12 | 5 | 3567 |
| Sex | Female | 804 | 345 | 185 | 137 | 103 | 72 | 49 | 43 | 15 | 18 | 7 | 6 | 1 | 3 | 0 | 1788 |
|  | Male | 586 | 311 | 188 | 179 | 122 | 134 | 80 | 60 | 42 | 22 | 15 | 14 | 12 | 9 | 5 | 1779 |
| IQ | High IQ | 464 | 208 | 101 | 85 | 67 | 56 | 29 | 29 | 13 | 12 | 8 | 5 | 4 | 3 | 1 | 1085 |
|  | Low IQ | 417 | 226 | 124 | 118 | 87 | 74 | 58 | 42 | 24 | 20 | 8 | 11 | 4 | 7 | 3 | 1223 |
|  | Mid IQ | 509 | 222 | 148 | 113 | 71 | 76 | 42 | 32 | 20 | 8 | 6 | 4 | 5 | 2 | 1 | 1259 |
| Medication | Medicated | 8 | 5 | 12 | 12 | 26 | 18 | 23 | 27 | 21 | 11 | 10 | 7 | 6 | 8 | 3 | 197 |
|  | Unmedicated | 1382 | 651 | 361 | 304 | 199 | 188 | 106 | 76 | 36 | 29 | 12 | 13 | 7 | 4 | 2 | 3370 |

Note: IQ groups defined by sample tertiles: Low ( $< 17$ ), Mid ( $17 \leq IQ < 20$ ), and High ( $\geq 20$ ). Due to the discrete nature of the IQ scores, the resulting groups are not exactly equal in size. Medication indicates stimulant use at assessment.

We use linear regression to test how ADHD scores, Sex, IQ, and Medication predict four basic behavioral measures (Table J): mean Go success reaction time (GSRT), Stop signal reaction time (SSRT), Go success (GS) rate, and Stop success (SS) rate. Note that SSRT estimates derived from the standard race model may be biased because the ABCD task design can violate the model’s independence assumption. The results show several statistically significant, yet very subtle associations. Higher ADHD scores significantly predict lower GS rates (*p <* 0.001). Among the covariates, Sex predicts GSRT and SSRT (*p <* 0.01); IQ predicts GSRT, SSRT, and GS rate (*p <* 0.01); and Medication predicts SSRT, GS rate (*p <* 0.001), and SS rate (*p <* 0.05). Most importantly, across all models, these effects are extremely small (all Partial *R*^2^ *≤* 0.025 and *f* ^2^ *≤* 0.026).

**Table J.** Linear model results for model-agnostic behavioral metrics.

| Behavioral metrics | Coef. | SE | *p*-value | Partial $R^2$ | $f^2$ |
| --- | --- | --- | --- | --- | --- |
| **ADHD** |  |  |  |  |  |
| GSRT | 0.2521 | 0.4315 | 0.5591 | 0.0001 | 0.0001 |
| SSRT | 0.2133 | 0.3516 | 0.5440 | 0.0001 | 0.0001 |
| GS rate | -0.0009 | 0.0003 | 0.0003*** | 0.0037 | 0.0037 |
| SS rate | $< 0.0001$ | 0.0001 | 0.8196 | $< 0.0001$ | $< 0.0001$ |
| **Sex (Male)** |  |  |  |  |  |
| GSRT | -6.5722 | 2.0872 | 0.0017** | 0.0028 | 0.0028 |
| SSRT | -4.6760 | 1.7004 | 0.0060** | 0.0021 | 0.0021 |
| GS rate | 0.0016 | 0.0013 | 0.2092 | 0.0004 | 0.0004 |
| SS rate | -0.0002 | 0.0007 | 0.7160 | $< 0.0001$ | $< 0.0001$ |
| **IQ** |  |  |  |  |  |
| GSRT | -1.3156 | 0.2909 | $< 0.0001$*** | 0.0057 | 0.0057 |
| SSRT | -0.7162 | 0.2370 | 0.0025** | 0.0026 | 0.0026 |
| GS rate | 0.0017 | 0.0002 | $< 0.0001$*** | 0.0248 | 0.0255 |
| SS rate | -0.0001 | 0.0001 | 0.2406 | 0.0004 | 0.0004 |
| **Medication** |  |  |  |  |  |
| GSRT | 1.7456 | 4.9027 | 0.7218 | $< 0.0001$ | $< 0.0001$ |
| SSRT | -12.4613 | 3.9941 | 0.0018** | 0.0027 | 0.0027 |
| GS rate | 0.0117 | 0.0030 | $< 0.0001$*** | 0.0044 | 0.0044 |
| SS rate | 0.0038 | 0.0016 | 0.0168* | 0.0016 | 0.0016 |

Note: $N = 3,567$ . \* $p < .05$ , \*\* $p < .01$ , \*\*\* $p < .001$ .

